# Multiplex imaging of human induced pluripotent stem cell-derived neurons with CO-Detection by indEXing (CODEX) technology

**DOI:** 10.1101/2022.02.03.479039

**Authors:** Laurin Heinrich, Faria Zafar, C. Alejandra Torres, Jasmine Singh, Anum Khan, Max Yang Chen, Cassandra Hempel, Nadya Nikulina, Jonathan Mulholland, Oliver Braubach, Birgitt Schüle

**Affiliations:** Department of Pathology, Stanford University School of Medicine, Stanford, CA, USA; Akoya Biosciences, Menlo Park, CA, USA; Cell Sciences Imaging Facility (CSIF), Beckman Center, Stanford University, Stanford, CA, USA

**Keywords:** Human-induced pluripotent stem cells, neuronal differentiation, iPSC, co-detection by indexing, CODEX, ultra-high multiplexed imaging, neuronal phenotyping

## Abstract

**Background:** Human induced pluripotent stem cell (iPSC) models have been hailed as a breakthrough for understanding disease and developing new therapeutics. The major advantage of iPSC-derived neurons is that they carry the genetic background of the donor, and as such could be more predictive for clinical translation. However, the development of these cell models is time-consuming and expensive and it is thus critical to maximize biomarker readout from every model that is developed. One option is to use a highly multiplexed biomarker imaging assay, like CO-Detection by indEXing (CODEX), which allows detection of 50+ targets *in situ* at single-cell resolution.

**New Method:** This paper describes the development of CODEX in neuronal cell cultures derived from human iPSCs.

**Results:** We differentiated human iPSCs into mixed neuronal and glial cultures on glass coverslips. We then developed and optimized a panel of 21 antibodies to phenotype iPSC-derived neuronal subtypes of cortical, dopaminergic, and striatal neurons, as well as astrocytes, and pre-and postsynaptic proteins.

**Comparison with existing methods:** Compared to standard immunocytochemistry, CODEX oligoconjugated fluorophores circumvent antibody host interactions and allow for highly customized multiplexing.

**Conclusion:** We show that CODEX can be applied to iPSC neuronal cultures and developed fixation and staining protocols for the neurons to sustain the multiple wash-stain cycles of the technology. Furthermore, we demonstrate both cellular and subcellular resolution imaging of multiplexed biomarkers in the same samples. CODEX is a powerful technique that complements other single-cell omics technologies for in-depth phenotype analysis.

**Graphical abstract:** Graphical abstract legend:
CODEX^®^ Multiplex Imaging in human iPSC neurons
[**A-D**] Schematic drawings of the tools and steps used for Co-Detection by indEXing (CODEX) imaging. [**A**] Target-specific antibodies are conjugated to unique DNA oligonucleotide barcodes. Fluorescent reporter (excitation wavelength at 488 nm, 550 nm, or 647 nm/Cy5) carrying the complementary DNA (to the barcode) enables barcode-specific binding of the reporter to the antibody and detection by fluorescence microscopy. [**B**] Neuronal cell cultures are prepared for the CODEX staining and imaging by several fixation steps with different PFA concentrations followed by incubation with 100% acetone. Residual acetone is removed by drying the sample. After rehydration with PBS, autofluorescence is quenched by exposure to broad-spectrum LED light. Following a pre-staining fixation step, the sample is incubated with a mix of all conjugated primary antibodies. Excessive, unbound antibodies are removed by a washing step, leaving only the bound antibodies followed by a final post-staining fixation. [**C**] The CODEX Instrument Manager performs the multicycle run and controls the microscope software for automated addition of reporters, imaging, and washing of the samples (pre-stained with primary antibodies) to remove reporters from each cycle. After imaging, bound reporters are removed without damaging the tissue using a solvent, and the next set of reporters (conjugated to different barcodes) are added. [**D**] CODEX^®^ Processor processes raw files and performs stitching, deconvolution, background subtraction, and cell segmentation. The processed images can be viewed and analyzed with the CODEX^®^ Multiplex Analysis Viewer (MAV) plugin using Fiji software.

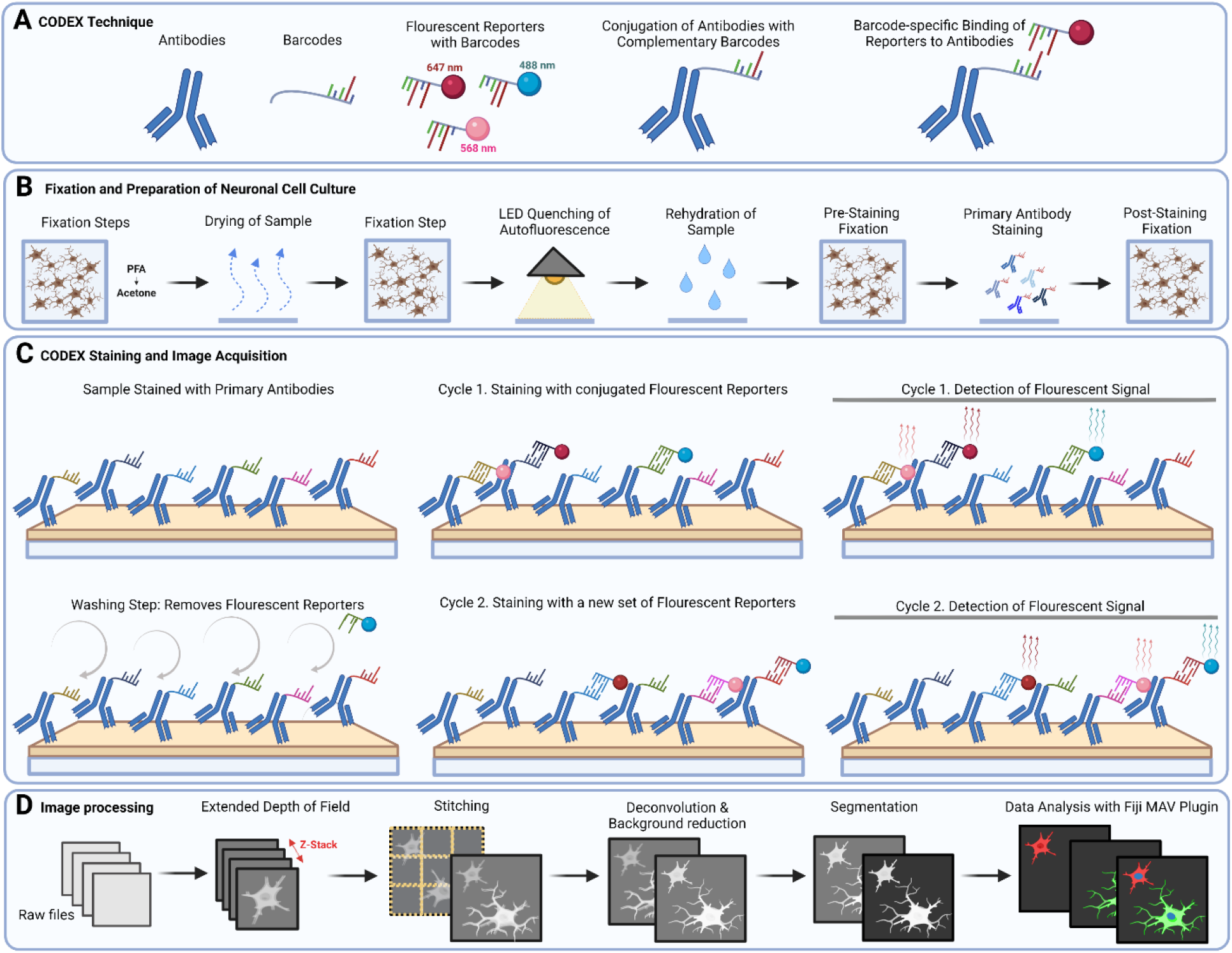

## 1. Introduction

Human induced pluripotent stem cell (iPSC) models are effective tools for studying human disease and developing therapeutic approaches for personalized medicine (Schüle et al., 2009). iPSCs can be generated from human somatic cells by induction of specific reprogramming factors and then further differentiated into various cell types, including neurons (Chang et al., 2019). A major advantage of human iPSC models is that they carry the human genetic background of the donor. Generating iPSCs from patients allows studying how genetic alterations lead to cellular and molecular changes, promoting the discovery of disease mechanisms and novel therapeutic targets. However, iPSC models have several challenges. Differentiation protocols are labor-intensive, costly, lengthy, and can be variable between experiments and clonal lines. It is therefore critical to maximizing the readouts of these *in vitro* models.

Developing novel multiplex technologies will increase the input-output ratio for the characterization and phenotyping of cellular subtypes and precursor cells of iPSC-derived cultures. CODEX^®^ imaging has been shown to multiplex up to ~56 markers per sample (Schürch et al., 2020), enabling extensive and deep characterization of samples. CODEX^®^ is based on the principle that specific antibodies are conjugated to unique oligonucleotide barcodes, which are recognized by a complementary CODEX^®^ reporter. The CODEX^®^ reporter has an oligo-conjugated fluorescent tag, which can be detected by conventional epifluorescence microscopy. Capitalizing on the reversible binding between the reporter to the antibody, already imaged reporters can be removed and washed out. Removing the reporters by a solvent-based washing procedure ensures the preservation of the tissue morphology and the primary antibody staining (Black et al., 2021). Staining, detection, removal can be repeated in multiple cycles.

While CODEX^®^ has been primarily employed for multiplex imaging of intact tissue sections, its application has not yet been demonstrated for iPSC-derived neuronal cultures. Of particular concern are the repeated wash and reporter readout steps that are required for CODEX imaging, given that these could pose a challenge and result in loss of precious cells during the experimental procedure. The protocol that we have developed overcomes this concern and enables deep characterization of iPSC-derived neurons with the CODEX^®^ technique and with virtually no loss of cellular substrate. Moreover, by imaging the same cell culture at both low and high magnification, we demonstrate that it is possible to generate highly multiplexed readouts at subcellular resolution. Although we have applied our approach to investigate and characterize iPSC-derived neurons, our work can also serve as a template for CODEX^®^ imaging of other cell-based model systems.

## 2. Material and Methods

### 2.1 iPSC maintenance

Human iPSCs were stored in Bambanker media in liquid nitrogen. Before thawing the iPSCs, one well of a 6-well plate was coated with Matrigel diluted 1:80 in Knockout Dulbecco’s modified Eagle’s (KO-DMEM) for 1 h at 37°C. Frozen vial with cells was quickly thawed for 2 min in a 37°C water bath. The cell suspension was transferred to 5 ml of Stemflex media and pelleted by centrifugation at 1000 rpm for 3 min. The supernatant was removed by aspiration and the cell pellet was resuspended in 1 ml of Stemflex media containing 1 μM of Thiazovivin (THZ). After removing the Matrigel solution from the coated well and adding 1 ml of Stemflex media containing 1 μM THZ, the cell suspension was added and mixed by gently shaking the plate. 48 h after plating, the media was changed to 3 ml Stemflex media without THZ.

Subsequently, media was changed every 48 hr. Once cells reached confluency they were passaged non-enzymatically with ReLeSR passaging solution. For passaging, media was aspirated and cells were washed once with 1ml phosphate-buffered saline (PBS) without Ca^2+^ and Mg^2+^ and then incubated with 1 ml ReLeSR for 4-6 min at room temperature (RT). After carefully aspirating off the ReleseR, cells were detached by gently pipetting with 1 ml Stemflex media containing 1 μM THZ.

Details on vendors and catalog numbers are listed in Supplemental Table 6 (Materials and Reagents).

### 2.2 Neuronal differentiation of iPSCs

#### 2.2.1 Neuronal differentiation by chemical induction

To generate iPSC-derived cortical, striatal, and dopaminergic neurons, we adapted three different protocols (Calatayud et al., 2021). First, forebrain PAX6-positive neural progenitor cells (NPCs) pre-patterned to derive into cortical neurons were generated from iPSCs by chemical induction using an adapted protocol from Shi *et al*., 2012 (Shi et al., 2012). To increase the outcome of PAX6 positive cells, we applied dual SMAD-inhibition by LDN/SB431542 instead of noggin/SB431542 (Surmacz et al., 2012). We further applied wingless/integrated (Wnt) pathway inhibition by exposure to IWP-2 (1 μM) during the first 4 days (Moya et al., 2014). Second, lateral ganglionic eminence (LGE) NPCs pre-patterned to derive into GABAergic medium-sized spiny neurons (MSNs) were generated from iPSCs by chemical induction using a modified protocol of Arber *et al*., 2015 (Arber et al., 2015). Exposure to SB431542 was reduced to 6 days (Day 0 to Day 6), while we started to supplement Activin A on day 8. We further adapted the LDN concentration to 500 nM and included IWP-2 (1 μM) during the first 4 days (Day 0 to Day 4). Third, floorplate NPCs pre-patterned to derive into dopaminergic neurons were generated from iPSCs by chemical induction using a modified dual SMAD-inhibition protocol as described by Kriks *et al*.,2011 (Kriks et al., 2011). The concentration of LDN was adapted (500 nM) and the timepoint of FGF8 exposure was optimized starting subsequently to sonic hedgehog/purmorphamine exposure from day 7.

For all three protocols, human KOLF2.1 iPSCs were first pre-patterned to NPCs. For generating NPCs, human iPSCs were passaged with ReLeSR and resuspended in a 1:1 mixture of Stemflex and KnockOut Serum Replacement (KSR) media (15% KSR, 1X Penicillin Streptomycin, 1X GlutaMAX, 1X non-essential amino acids (NEAA), 0.1 mM beta-mercaptoethanol in KO-DMEM/F12) containing 1 μM THZ. Cells were plated at a concentration of 300,000 cells/cm^2^ on freshly-coated Matrigel plates (1:50 in KO-DMEM).

For differentiation of NPCs into the respective neuron types, NPCs were plated onto primary human astrocytes grown on poly-L-lysine (PLL)/laminin-coated coverslips (Section 2.4.2). One day prior, 300,000 cells of 2 mM Ara-C-inactivated primary human astrocytes were thawed and plated in 400 μl Astrocyte media onto the CODEX coverslip (22×22 mm) only and allowed to attach for 10 min. Then 1 ml of astrocyte media was carefully added to the 6-well plate.

The next day, NPCs were passaged using ReLeSR and collected in N2B27 (Neurobasal medium: DMEM/F12 (1:1), 0.5X B27 without vitamin A (Vit.A), 0.5X N2, 1X PenStrep, 1X GlutaMAX, 1X NEAA) media containing 1 μM THZ. By careful pipetting NPCs were plated onto the coverslip with the pre-seeded primary astrocytes and allowed to attach for 10 min. Then 2 ml of N2B27 media containing 1 μM THZ was gently added to the well. To induce differentiation of the NPC the media was changed to terminal differentiation media (1X B27 without Vit. A, 1X Glutamax, 1X Pen Strep, 1X NEAA, 0.5 mM dbcAMP, 0.2 mM L-Ascorbic Acid, 1 μM DAPT, 100 nM SR11237, freshly added: BDNF 10 ng/ml, GDNF 10 ng/ml, and 0.5% fetal bovine serum (FBS) in Neurobasal A) the next day. For the first 14 days, half of the media was changed every 3 days and 1 μg/ml of mouse laminin was added to the media every second time of media change.

After day 14, half of the media was changed every 7 days and 1 μg/ml of mouse laminin was added to the media every second time of media change. After growing the culture in terminal differentiation media for 7 days, Culture One treatment (1X) was added to the media for three weeks to stop the growth of undifferentiated progenitor cells.

#### 2.2.2 Generation of Ngn2-induced cortical neurons

The Neurogenin 2 (Ngn2) iPSC line (name: WTC11_G3) was differentiated into cortical neurons with a two-step differentiation protocol (pre-differentiation and maturation) (Wang et al., 2017). For pre-differentiation (Days −3 to −1), iPSCs were seeded (Day −3) in KO-DMEM/F12 medium containing 1X N2 supplement, 1X NEAA, 1x GlutaMAX, 0.2 mg/mL mouse laminin, 10 ng/mL BDNF, 10 ng/mL neurotrophin-3 (NT3), 2 mg/mL doxycycline and 1 μM THZ at a density of 200,000 cells/cm^2^ in a 6-well plate coated with Matrigel. The medium (without THZ) was then changed daily for two more days, and THZ was removed from day-2. 300,000 cells of 2 mM Ara-C-inactivated primary human astrocytes were thawed and plated in 400 μl astrocyte media onto the PLL/laminin-coated coverslip only and allowed to attach for 10 min. Then 1 ml of astrocyte media with 0.5% FBS was carefully added to the 6-well plate. For maturation at day 0, pre-differentiated precursor cells were dissociated using ReLeSR, and plated at 150,000 cells/cm^2^ density on astrocyte-feeder plates in maturation medium containing 50% DMEM/F12, 50% Neurobasal-A medium, 0.5X B27 supplement, 0.5X N2 supplement, 1X GlutaMax, 1X NEAA, 1 mg/mL mouse laminin, 10 ng/mL BDNF, 10 ng/mL NT3, 0.5% FBS and 1 μM THZ. Half of the medium was replaced on Day 1, Day 3, Day 7, and again on Day 14. Thereafter, half of the medium was replaced weekly until the cells were fixed for staining.

### 2.3 Characterization and validation of iPSC-derived neurons using Q-PCR by SYBR Green multi-well array

iPSCs and iPSC-derived neurons were characterized and validated using a previously established 96 -well SYBR Green qPCR expression array (Srinivasaraghavan et al., 2022). A confluent well from a 12-well plate of iPSCs (^~^2,500,000 cells) or one 6-well of iPSC-derived neurons (^~^500,000 cells) were dissociated with ReLesR. iPSCs were pelleted for 5 min by centrifugation (5,000 g at 4°C), washed with 1 ml of PBS again, followed by another 5 min centrifugation step (5,000 g at 4°C). RNA was then extracted using the Purelink^™^ RNA Mini Kit. To increase RNA yield, differentiated neurons were in some cases treated with lysis buffer directly on the plate without dissociating and pelleting cells first. Cells were washed with 1 ml PBS twice, then 300 μL of lysis buffer containing 3 μL of 2-mercaptoethanol was added directly to the well for 1-2 min at RT. The cell lysis was directly transferred to the homogenization tube. All subsequent steps of the Purelink^™^ RNA extraction protocol were identical for iPSCs and neurons. The volume of lysis buffer was adjusted based on cell numbers, according to the Purelink^™^ kit protocol. To eliminate contamination with genomic DNA, 1 μg of RNA were treated with DNase I for 15 min at RT followed by 65°C inactivation for 3 min and then directly used for cDNA reverse transcription. using the High-Capacity cDNA Reverse Transcription kit. Negative control was generated with nuclease-free water instead of RNA, and a reverse transcriptase control was generated with RNA but no reverse transcriptase. Afterward, 80 μl of nuclease-free water was added to each cDNA tube for a final concentration of 10 ng/μL and a final volume of 100 μL before storing at −20°C.

For the SYBR Green array, both forward and reverse primers (10 μM in nuclease-free water) of the target gene were mixed in a stock solution and coated at 0.3 μL (final concentration of 300 nM each primer) into a 384-well of an optical PCR plate in triplicates. On every plate, a negative control containing GADPH primers and the negative control sample (mentioned above) was included. After finishing the pre-coating of all primers for all genes of interest, the 384 well-plate was centrifuged at 2,000 rpm for 2 min. Plates were then placed in a box or drawer and pre-coated primers were left to dry overnight at RT. The SYBR Green array reaction was run in a QuantStudio 6 Flex with cycling conditions as indicated in Supplemental Tables 8 and 9. A reaction mix containing 2.5 μL of PowerUp SYBR Green Master Mix, 1.5 μL of nuclease-free water, and 1 μL of sample cDNA at 10 ng/μL (final cDNA concentration of 2 ng/μL) was prepared per well. After pipetting the reaction mix to the wells with the pre-coated primers, the plate was sealed with optical adhesive film and centrifuged at 2,000 rpm for 2 min. The PCR conditions in Supplemental Table 8 are run first to amplify cDNA, while the conditions in Table 9 were run at the end to generate a melt curve from the resulting PCR product.

To quantify relative gene expression among different cell types, the fold change was calculated from cycle threshold (Ct) values. Mean Ct values for each gene and sample were calculated among each triplicate. ΔCt was then calculated using *ΔCt = mean Ct_target gene_ – mean Ct_housekeeping gene (GAPDH)_*. ΔΔCt was then calculated using *ΔΔCt = ΔCt_treated cell (neuron)_ - ΔCt_untreated cell (iPSC)_*. Finally, fold change was calculated using *Fold change = 2^−ΔΔCt^*. ΔCt represents the difference in gene expression between the gene of interest and the housekeeping gene. ΔΔCt normalizes the relative gene expression to an untreated control sample. For this study, the respective untreated iPSC line was used as a reference sample. Markers specific for various neuron types, pluripotency, neuronal maturation, and stress were used. Supplemental Table 7 enlists all primers and Supplemental Table 6 shows details on reagents and materials.

### 2.4 CODEX^®^ imaging

#### 2.4.1 Preparation of coverslips for neuronal cell culture

For CODEX^®^, cells were plated and differentiated on specific 22×22 mm glass coverslips (Electron Microscopy Sciences, 72204-01) that fit the frame of the PhenoCycler instrument. To enable attachment and long-term culture of cells, the coverslips were treated with poly-L-lysine for 12 h at RT, washed once with double-distilled water (ddH2O), and dried for 1 h at RT before single coverslips were placed in individual wells of a 6-well plate and sterilized under UV-light for 30 min. Afterward, the coverslips were coated with 10 μg/ml mouse laminin diluted in sterile ddH2O for 2 h at RT. The laminin was taken off the coverslip and allowed to dry for 2 min at RT before cells were plated (Sections 2.2.1 and 2.2.2).

#### 2.4.2 Pre-staining fixation and autofluorescence quenching of neurons on the coverslip

Coverslips with mixed iPSC-derived cortical, striatal, and dopaminergic neurons for 60-70 days *in vitro* (DIV) and cortical neurons for 45DIV were processed for initial fixation (Zafar et al., 2022). 1.5 ml media (of 3 ml total media) was taken off the well and 1.5 ml 4% paraformaldehyde (PFA)/PBS solution was added slowly to the well and incubated at RT for 10 min. Next, media/PFA solution mix was carefully aspirated from the well with a micro pipettor, and 2 ml 4% PFA/PBS solution was added at RT for 5 min.

Afterward, cells were washed with PBS twice. Next, coverslips were placed (cells facing up) in a box containing Drierite absorbent beads at RT for 2 min. Cells were fixed again by placing the coverslip with forceps into 10 ml of 100% acetone in a 50 ml beaker at RT for 5 min (Figure 1 Initial Fixation Steps). To remove acetone by evaporation, coverslips were taken from the beaker with forceps and placed onto Drierite beads for 2 min at RT again. Cells were finally fixed with 1.6% PFA in 5 ml Akoya of Hydration Buffer at RT for 10 min and washed with 5 ml PBS three times. To quench auto-fluorescence, the coverslips were then placed in fresh PBS in a well of a 6-well plate and exposed to a broad-spectrum LED light (DCenta A4 Ultra Bright LED Light Box Tracing Pad 25000Lux) for 45 min at RT. Then sample was transferred to fresh PBS and exposed to broad-spectrum LED light again for 45 min at RT. Finally, samples were equilibrated by placing them in Akoya Hydration buffer twice for 2 min each before staining.

**Figure 1:**
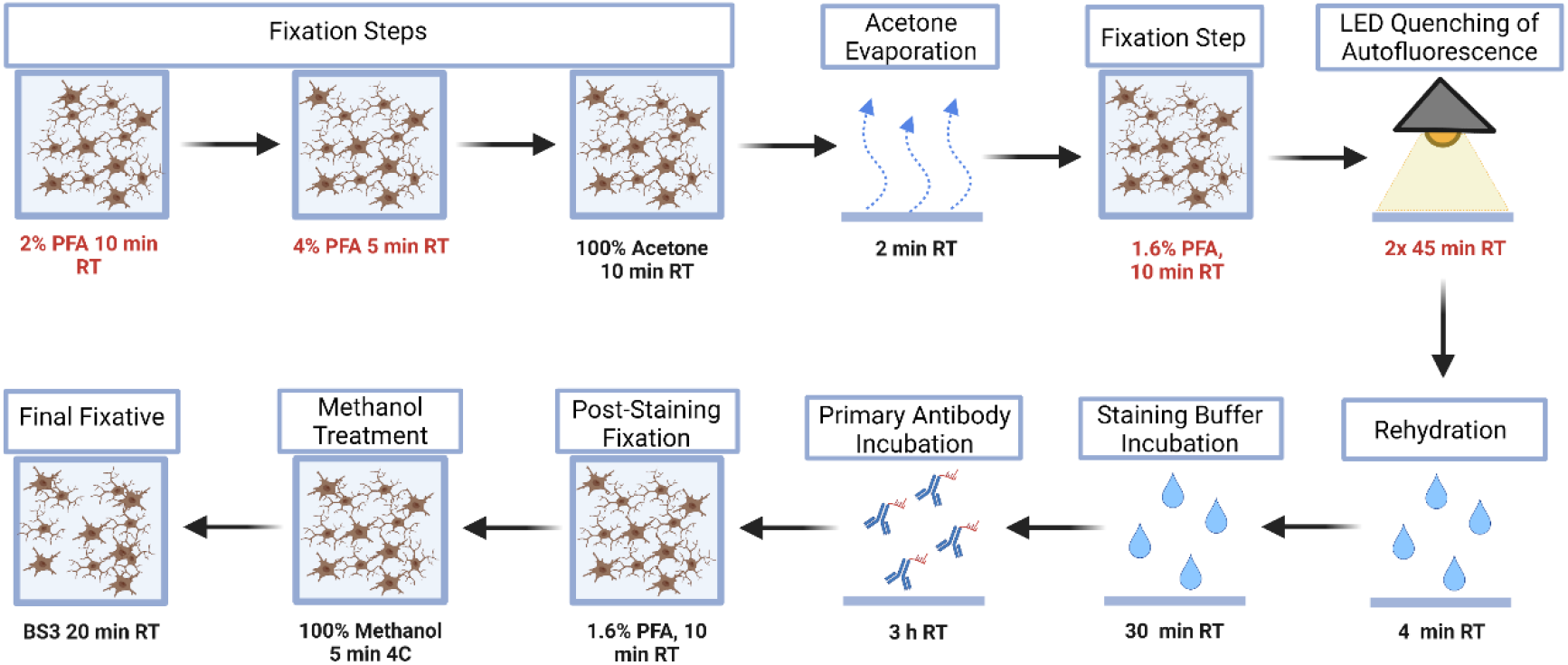
Workflow for the fixation and staining of iPSC-derived neurons for multiplex imaging with CODEX. The protocol for CODEX multiplex tissue imaging was adapted (modified steps are marked in red) for iPSC-derived neuronal in vitro cultures. iPSC-derived neurons cultured on a 22×22 mm coverslip are fixed with 2% PFA for 10 min (1:1 mixture of media and 4% PFA), with 4% PFA for 5 min, and then with 100% acetone for 10 min at room temperature (RT). For evaporation of acetone, the coverslip is placed on Drierite beads at RT for 2 min and then fixed again with 1.6% PFA in Hydration buffer for 2 min at RT. Autofluorescence is quenched with LED light twice for 45 min in PBS at RT. Subsequently, samples are rehydrated in Hydration buffer twice at RT for 2 min each and then incubated in Staining buffer for 30 min. Samples are stained with primary antibodies diluted in Staining buffer at RT for 3 h and fixed again with 1.6% PFA for 10 min at RT, with 100% Methanol for 5 min at 4°C, and with BS3 substrate for 20 min at RT.

**Figure 2:**
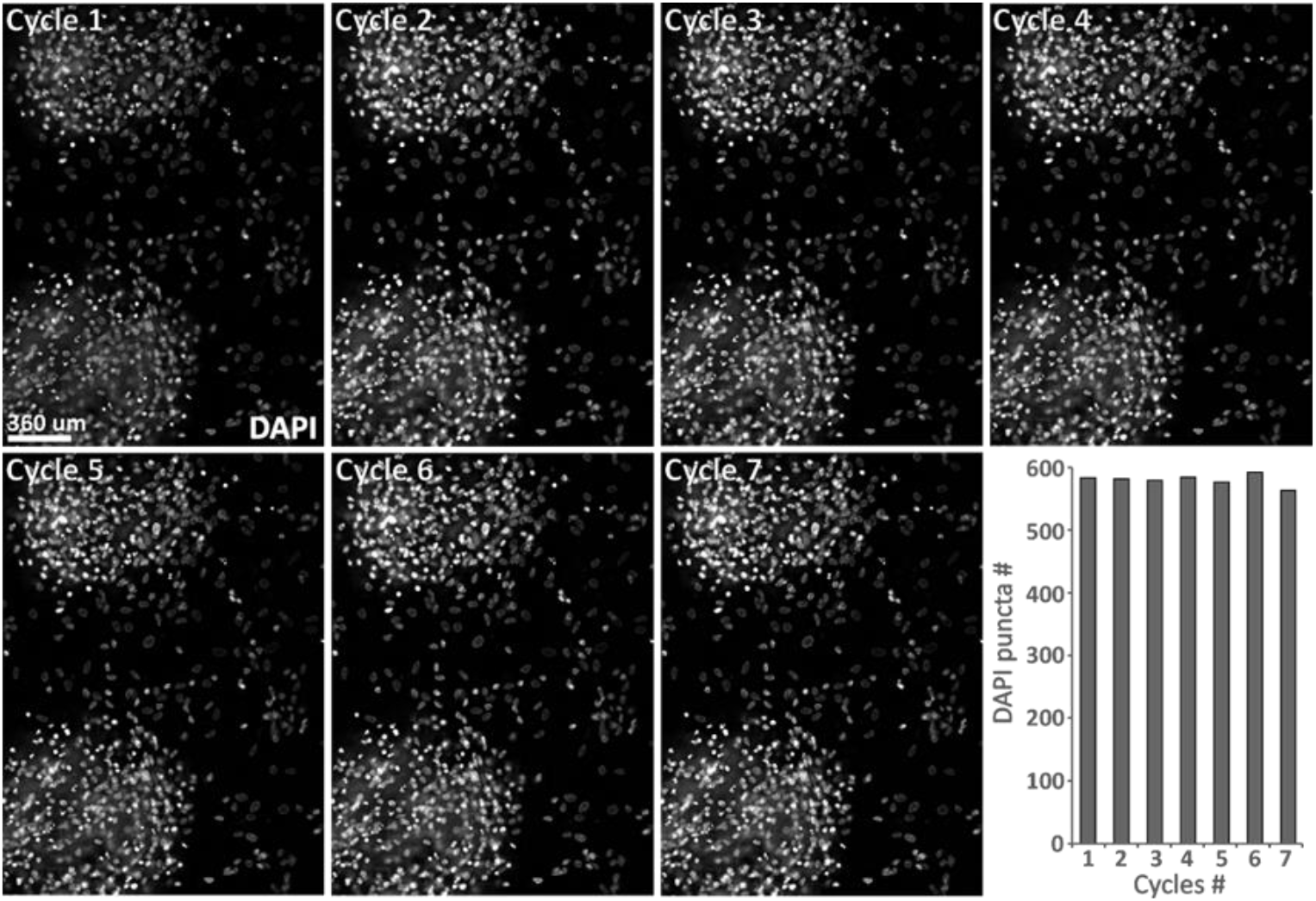
No overt cell loss between CODEX imaging cycles. Human KOLF2.1 iPSC-derived neural progenitors were co-cultured with primary human astrocytes and differentiated to neurons (60DIV). Codex imaging was performed for seven cycles. DAPI puncta were counted using ImageJ software to test for loss of cells between the imaging cycles.

#### 2.4.3 Primary antibody staining and post-staining fixation

Before starting the primary antibody staining, cells on the coverslips were equilibrated in Akoya Staining Buffer at RT for 30 min. The primary antibody cocktail was prepared in blocking buffer. Details regarding primary antibody clones, vendors, catalog numbers, barcode assignments, and concentrations are listed in Supplemental Tables 1-5. Cells on coverslips were incubated with a primary antibody/blocking buffer cocktail in a humidity chamber at RT for 3 h or overnight at 4°C (Figure 1). After incubation, coverslips were washed with Staining Buffer twice at RT for 2 min each. For post-fixation, coverslips (primary antibody incubated cells) were fixed with 1.6% PFA/Storage Buffer mix at RT for 10 min. Then, coverslips were washed with PBS three times at RT for 2 min each. Next, the coverslips were incubated with ice-cold 100% methanol on ice for 5 min. Coverslips were washed again with PBS, three times at RT for 2 min each. Final fixation was done using a 1:50 bis(sulfosuccinimidyl)suberate (BS3) to PBS mix at RT inside the humidity chamber for 20 min. Coverslips were washed with PBS three times at RT for 2 min each. At this point, coverslips can be stored for 5 days in storage buffer at 4°C. Laboratory safety and chemical waste guidelines were followed throughout all steps of this protocol.

#### 2.4.4 Imaging and raw-data processing

All imaging experiments were performed with a PhenoCycler^™^ platform connected to a Keyence BZ-X800 epifluorescence microscope. Before imaging, CODEX reporter 96-well plates were prepared using the cycle reporter configuration as outlined in the CODEX user manual (Akoya Biosciences, 2021a). Imaging was performed using a Keyence BZ-X800 epifluorescence microscope equipped with either a 20x objective (Nikon CFI Plan Apo 20x/0.75) or a 40x oil-immersion objective (Nikon, Plan Fluor, 1.30 NA). A large region was initially identified using the stitching option in BZ-X800 viewer software, and z-stacks were then acquired throughout this region with a total depth of 10 μm each. The multiplex cycles were set up using Akoya’s CODEX Instrument Manager (version 1.29.3.6). Acquired images were then processed, stitched, and segmented using the CODEX Processor set at default manufacturer values (Akoya Biosciences, 2021b). Final data were then viewed using the CODEX Multiplex Analysis Viewer (MAV) plugin (Akoya Biosciences, 2021c) for FIJI (ImageJ) (Schindelin et al., 2012).

More details on the CODEX fixation, staining, fluorophore-labeled reporter setup, and imaging can be found in the CODEX user manual (Akoya Biosciences, 2021a).

## 3. Results

### 3.1 Design of CODEX antibody panel

#### 3.1.1 Rationale for choosing markers in CODEX panel

We developed a CODEX panel consisting of 21 markers to characterize and phenotype iPSC-derived neurons (Figure 3–5). We included neuron-specific and glia markers that allowed us to distinguish cell types and different neuronal subtypes at various stages of maturity and/or disease and cell damage.

**Figure 3:**
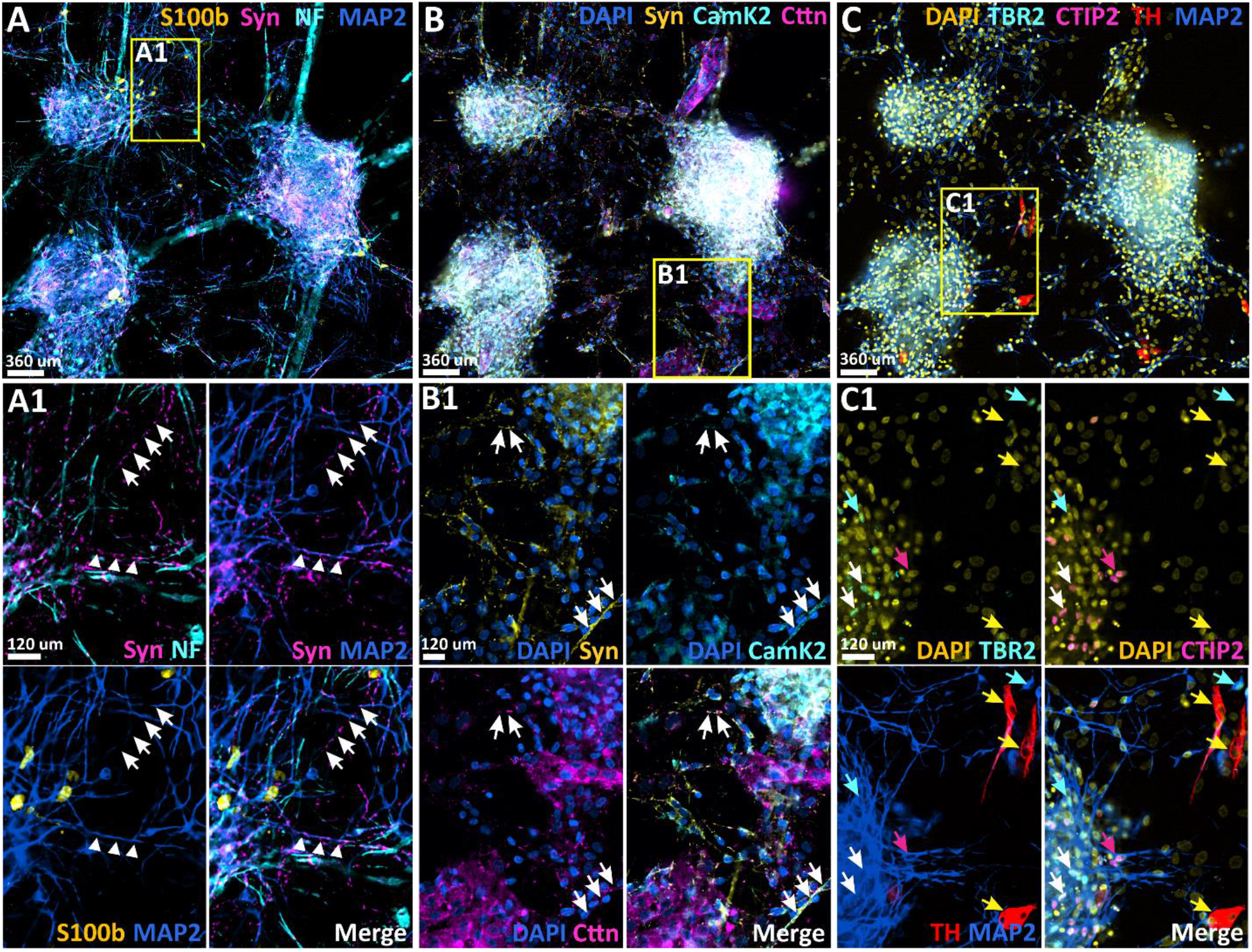
CODEX Multiplex Imaging of iPSC-derived Cortical and Dopaminergic neurons co-cultured with primary human Astrocytes (60DIV). Human KOLF2.1 iPSC-derived neural progenitor cells were co-cultured with primary human astrocytes and differentiated to neurons. [**A-C**] Nine different markers are shown to identify specific cell types and/or maturity levels of cells. Scale bars represent either 360 μm or 120 μm. [**A**] Representative image of S100β (yellow), synapsin 1 (Syn, magenta), neurofilament (NEFL, cyan), and MAP2 (blue) staining. [**A1**] Magnification of the selected area (yellow square in **A**) highlighting SYN co-localization with NEFL (white triangles) and MAP2 (white arrows). [**B**] Representative image of DAPI (yellow), TBR2 (cyan), CTIP2 (magenta), tyrosine hydroxylase (TH, red), and MAP2 (blue) staining. [**B1**] Magnification of selected area (yellow square in **B**) highlighting TBR2 co-localization with DAPI (cyan arrows), CTIP2 co-localization with DAPI (magenta arrows), as well as co-localization of CTIP2 with TBR2 (white arrows). [**C**] Representative image of DAPI (blue), synapsin (SYN, yellow), CAMK2 (cyan), and Gephyrin (Geph, magenta) staining. [**C1**] Magnification of selected area (yellow square in **C**) highlighting CAMK2 co-localization with SYN (white arrows).

**Figure 4:**
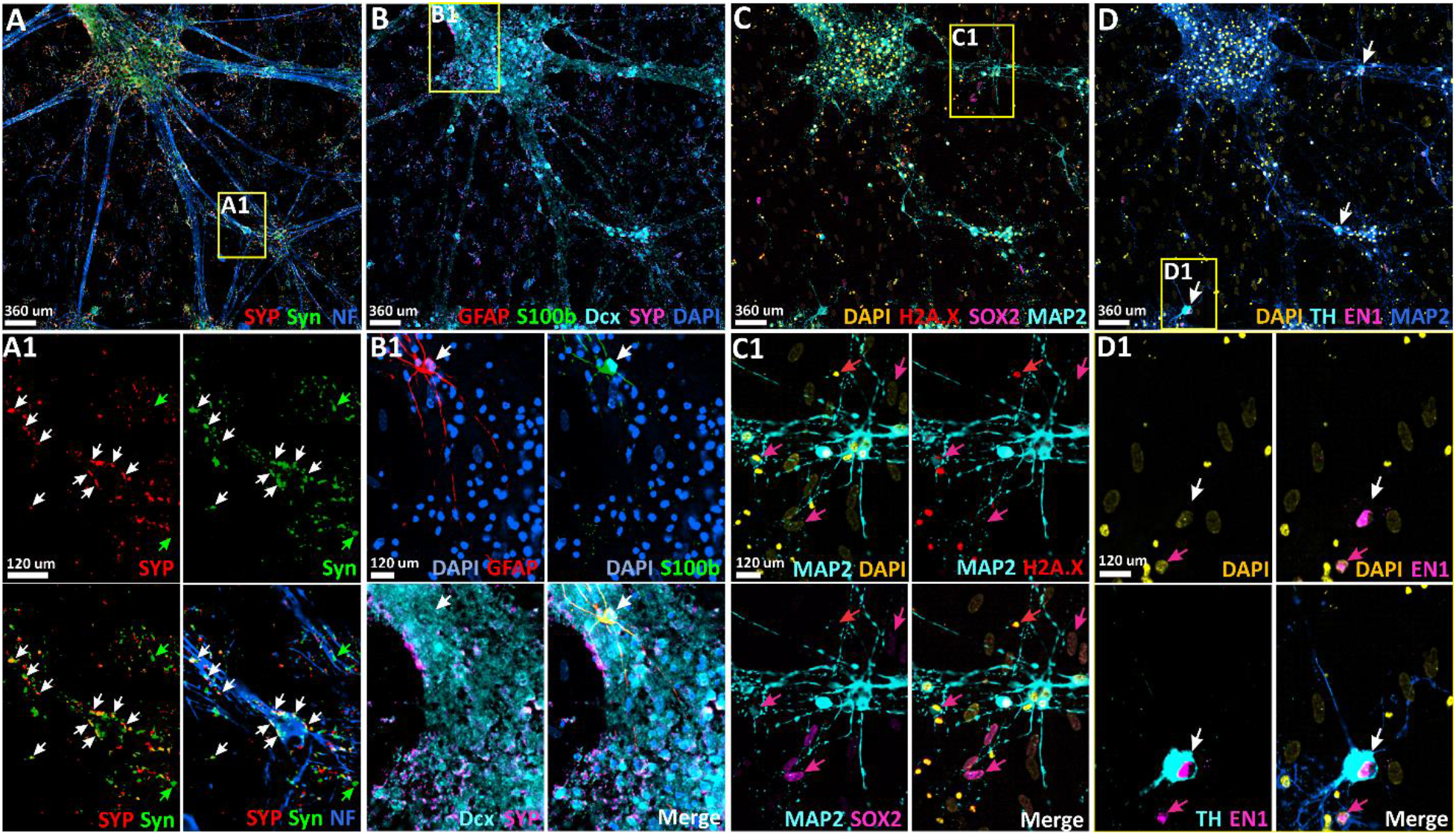
CODEX Imaging of iPSC-derived cortical neurons co-cultured with primary human astrocytes (45 DIV). Human Ngn2 iPSCs were co-cultured with primary human astrocytes and differentiated to neurons with cortical identity. [**A-D**] Codex imaging was performed for 11 different markers to identify specific cell types and/or maturity. Scale bars represent either 360 μm or 120 μm. [**A**] Representative image of synaptophysin (SYP, red), synapsin 1 (SYN, green), and neurofilament (NF, blue) staining. [**A1**] Magnification of the selected area (yellow square) highlighting SYP co-localization with SYN (white arrows), SYP and SYN co-localization with NEFL (white arrows), and SYN signal not co-localizing with SYP (green arrows). [**B**] Representative image of GFAP (red), S100B (green), doublecortin (DCX, cyan), SYP (magenta), and DAPI (blue) staining. [**B1**] Magnification of selected area (yellow square) highlighting GFAP co-localization with S100B (white arrow), DCX patterning with SYP. [**C**] Representative image of DAPI (yellow), H2A.X (red), SOX2 (magenta), and MAP2 (cyan) staining. [**C1**] Magnification of selected area (yellow square) highlighting H2A.X co-localization with DAPI (red arrows) and SOX2 co-localization with DAPI (magenta arrows). [**D**] Representative image of DAPI (yellow), tyrosine hydroxylase (TH, cyan), engrailed 1 (EN1, magenta), and MAP2 (blue) staining. White arrows highlight TH positive cells. [**D1**] Magnification of the selected area (yellow square) highlighting TH co-localization with EN1 and MAP2 (white arrows), and EN1 signal not co-localizing with TH positive cells (magenta arrows).

**Figure 5:**
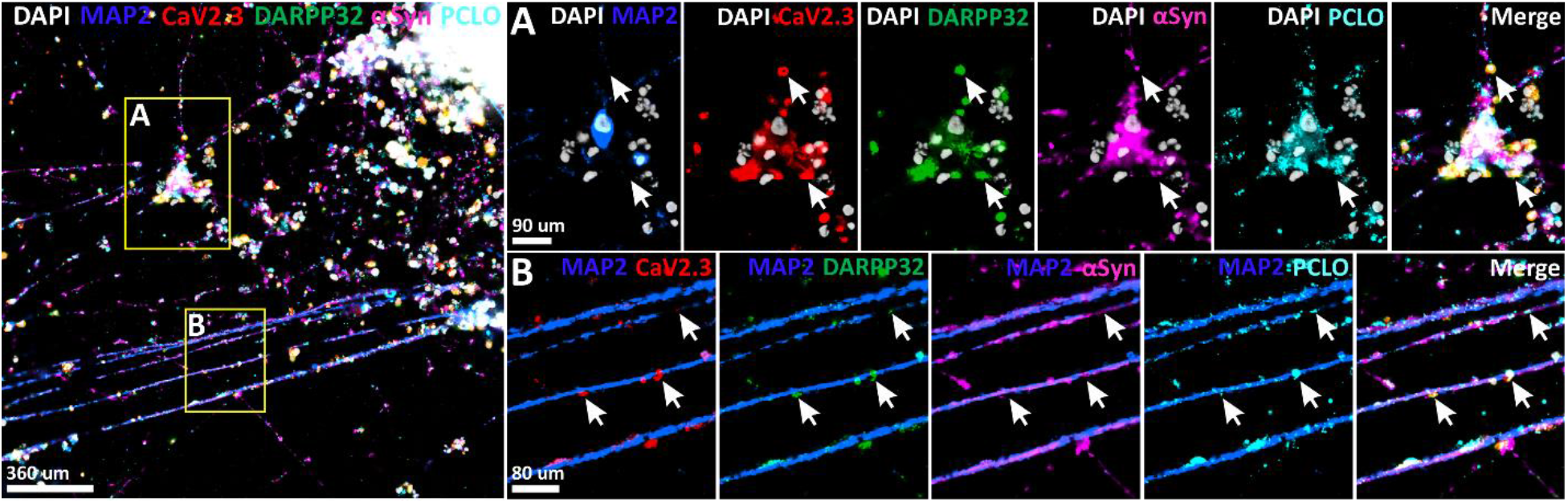
CODEX Imaging of iPSC-derived neurons (70DIV) using a 40x oil-immersion objective. Human KOLF2.1 iPSC-derived neural progenitor cells were differentiated into neurons. Codex imaging was conducted for 5 different markers using a 40x oil-immersion lens. Scale bars represent either 360 μm, 90 μm, or 80 μm. [**A, B insets**] DAPI (white), MAP2 (blue), CAV2.3 (red), DARPP32 (green), alpha-synuclein (aSyn, magenta), and piccolo (PCLO, cyan) staining.

For the identification of neurons, we optimized antibodies against the neuron-specific cytoskeletal proteins neurofilament light subunit (NEFL, Figure 3A and Figure 4A) (Gafson et al., 2020) and the microtubule-associated protein 2 (MAP2 Figure 3A and Figure 4C-D) (Cassimeris and Spittle, 2001). While neurofilaments are strongly enriched in axons, where they determine radial growth and the axon diameter (Gafson et al., 2020), MAP2 labels primarily dendritic structures as it is involved in dendrite growth and stabilization. The *MAP2* gene is further known to encode three different protein isoforms that are indicators for juvenile (MAP2c) and mature neurons (MAP2a and MAP2b) (Cassimeris and Spittle, 2001). To detect mature neurons, we used a MAP2a-specific antibody.

As synapses are major indicators of neuronal function and connectivity, we established antibodies against the presynaptic proteins synapsin 1 (SYN, Figure 3A-B and Figure 4A) and synaptophysin (SYP, Figure 4A) (Ashery et al., 2013). The neuron-specific phosphoprotein SYN is an early neuronal marker already present during neurogenesis where it is involved in axon elongation and synaptogenesis. SYN also plays a role at the presynaptic terminals of mature neurons where it binds to the cytoplasmatic side of synaptic vesicles and modulates synaptic activity (Mirza and Zahid, 2018). By contrast, the vesicular transmembrane protein SYP is a synaptic marker that appears later during development, where it is involved in the activity-dependent synapse formation and trafficking of synaptic vesicles to presynaptic terminals (White and Stowell, 2021). We also included markers for proteins important in neuronal development to further verify the neuronal identity of our iPSC-derived neurons, including the calcium/calmodulin-dependent protein kinase II (CAMK2 Figure 3B) (Kool et al., 2019; Wang et al., 2013), the actin-binding protein cortactin (CTTN, Figure 3B) (Cornelius et al., 2021), and the microtubule-associated protein doublecortin (DCX, Figure 4B) (Ayanlaja et al., 2017).

To investigate disease-relevant phenotypes in patient versus control iPSC-derived neurons, it is important to be able to generate and identify disease-relevant neuronal subtypes. We added a range of antibodies that can be used for classifying cortical, striatal GABAergic, and dopaminergic neurons. We included the transcription factor T-box transcription factor 2 (TBR2, Eomes (Eomesodermin), Figure 3C) and chicken ovalbumin upstream promoter-transcription factor-interacting protein 2 (CTIP2, BCL11B (BAF chromatin remodeling complex subunit), or Rit-1β, Figure 3C) as markers for excitatory cortical neurons. While TBR2 is involved in the cortical patterning of intermediate progenitors in the developing cerebral cortex (Hevner, 2019), CTIP2 is involved in the late development of cortical deep-layer projection neurons. Nevertheless, CTIP2 can also be used as a marker for GABAergic neurons, specifically medium-sized spiny neurons in the striatum, where it was shown to be most highly expressed (Arlotta et al., 2008). As markers for dopaminergic neurons, we added antibodies specific for tyrosine hydroxylase (TH, Figure 3C and Figure 4D) and engrailed homeobox 1 (EN1, Figure 4D). TH shows abundant expression in dopaminergic neurons where it catalyzes the rate-limiting step in the conversion of tyrosine to dopamine (Daubner et al., 2011). EN1 belongs to a group of transcription factors that controls the cell fate of midbrain dopaminergic neurons during development, but EN1 was also found to be expressed in mature midbrain dopaminergic neurons where it is critical for survival and maintenance (Rekaik et al., 2015).

Apart from neurons, we expected our culture to consist of astrocytes, as neuronal progenitor cells were co-cultured with primary human astrocytes, and undifferentiated stem and progenitor cells. Hence, we included the two markers S100 calcium-binding protein B (S100B; Figure 3A and Figure 4B) and the glial fibrillary acidic protein (GFAP; Figure 4B) that are highly expressed in astrocytes (Raponi et al., 2007). To detect undifferentiated cells, we stained for the SRY-box transcription factor 2 (SOX2; Figure 4C) a transcription factor found to be essential for the maintenance of pluripotency in embryonic stem cells (Holmes et al., 2020). As we were also interested in the viability of our cultured iPSC-derived neurons and stained against histone H2AX (H2A.X; Figure 4C) which is a specific and sensitive marker for DNA double-strand breaks and thereby an early indicator of neuronal damage (Crowe et al., 2011).

#### 3.1.2 Selection criteria for antibody clones

Our multiplexed imaging of neuronal *in vitro* cultures with the CODEX technology comprised a new protocol on a new substrate and thus required all of our antibodies to be procured, conjugated, and validated. We pre-selected antibodies based on the following criteria: (i) the conjugation process requires purified antibodies not containing the carrier protein bovine serum albumin (BSA), or other proteins (ii) a minimum antibody quantity of 50 mg is needed for conjugation, this concentration had to be confirmed with a nanodrop (iii) previously optimized antibodies for iPSCs and iPSC-neurons, (iv) monoclonal antibodies as they will have a higher specificity for their target proteins, (v) antibody clones already validated in immunohistochemistry or immunofluorescence in human brains, (vi) antibodies that have been validated using knockout mice or by positive and negative controls via Western immunoblotting.

Some antibodies were custom-ordered, processed to remove bovine serum albumin (BSA removal kit Abcam). The antibodies were conjugated to oligonucleotide barcodes according to instructions in the Akoya CODEX user manual (Akoya Biosciences, 2021a, chapter 4 Antibody Conjugation). Supplemental Tables 1-4 summarize antibodies and barcodes of the panel. Supplemental Table 5 summarizes the antibodies that were tested but failed or did not show positive staining in iPSC-derived neurons.

#### 3.1.3 Selection criteria for oligo barcodes and fluorescent reporters for the CODEX panel

Barcodes and reporters are an essential part of the CODEX technology. Our barcode-reporter selection was based on enhancing the signal-to-noise ratio and avoiding barcode-reporter repetition. We first selected and enlisted all desired antibodies along with their expected abundance and localization *in situ*.Then we performed an antibody search for the selected proteins and evaluated them based on vendor information and literature references. For untested antibodies, we first performed conventional immunocytochemistry to assess protein localization and abundance, optimal antibody concentration, and the exposure time required for imaging.

Each CODEX cycle consists of; a nuclear stain (405 nm wavelength/ blue) and three fluorophores AF488 (green), Atto550 (red), and Cy5 (far red), which needed to be assigned to individual reporters/barcodes per cycle. It is recommended to use reporters that emit in a higher wavelength like Cy5 for low abundance proteins because of high autofluorescence in the lower wavelength channels. Nevertheless, the final assignment of protein/reporter depends on the signal intensity obtained in the optimization process. Cy5 fluorophore signal is usually weaker than the one in 488 nm or 550 nm channel. If a marker shows a low signal in the Cy5 channel it can be moved to the 488 nm channel to increase the signal-to-noise ratio. Overall, the choice of fluorophore should be balanced between intrinsic autofluorescence and signal-to-noise ratios.

The final step in building the panel was the assembly of the multicycle CODEX experiment. Each pair of barcode/reporter had to match and not overlap between different antibodies. All desired antibodies were thus grouped in the three available channels (488 nm, 550 nm, and Cy5) and distributed to avoid “empty channels” in the experimental plan. These “empty channels” underwent the same image acquisition as the ones coupled with a reported therefore taking additional time and storage space. It is thus advantageous to design multiplex panels that use three antibodies per cycle.

### 3.2. Validation and quality control of barcode conjugated antibodies

Verification of the CODEX-barcode conjugation chemistry was performed by immunoblotting. Conjugated antibodies have a higher molecular weight due to the binding of the oligonucleotide to the antibody. Furthermore, successful conjugation was validated by manual immunostaining of test samples according to the workflow described in the method section. For validation purposes, imaging was performed manually and not conducted using the automated CODEX process. Test samples were counter-stained with diamidino-2-phenylindole (DAPI) and imaged using a digital fluorescent microscope. Validation of these conjugated antibodies was complete after staining was verified both using manual and multiplex CODEX runs. These two quality control steps allowed verification of the staining pattern, specificity, and signal-to-noise ratio and that its binding efficiency was not impacted by the presence of multiple antibodies. Lastly, as a sample quality control, we implemented a multi-well SYBR Green mRNA expression panel against proteins recognized by antibodies in our panel. As mRNA is required for synthesizing proteins, expression levels of mRNA in cells can be informative about the respective protein concentration (Alberts et al., 2002). Investigating the mRNA expression levels of specific genes will consequently help us determine whether, in the absence of a fluorescent signal, the antibody did not work, or the respective gene was not expressed in iPSC-derived neuronal cultures.

### 3.3. Image processing

For image processing, we applied the bioinformatics tools provided by Akoya. Segmentation proved to be challenging, as the processor’s segmentation algorithm considers cells to be predominantly circular and neurons have more complicated shapes with processes. Hence, new software or improved algorithm designed for segmenting neuronal cell types would need to be developed for future applications and CODEX imaging of iPSC-derived neuronal networks.

### 3.4. Improved protocol for the preparation, fixation, and incubation of antibodies of iPSC-derived neurons allows multiplex imaging by CODEX

A hurdle of staining and imaging neuronal cultures is the risk of losing cells even during standard washing and staining procedures as iPSC neurons have fragile processes, and because attachment of cells over a long period in culture is reduced. Therefore, iterative multiplex imaging applications such as CODEX, demand a specialized preparation and fixation protocol.

In this study, we optimized the protocol for multiplex tissue imaging and adapted it for neuronal cell cultures. Briefly, we found that additional fixation steps (Figure 1) increase the durability of iPSC-derived neuronal cultures and enable CODEX imaging without obvious cell loss. We found that pre-fixation of neuronal cultures first with 2% PFA for 5 min and second with 4% PFA for 10 min before acetone incubation enhanced the durability of neuronal networks during the subsequent staining procedures. 100% acetone treatment was performed to dehydrate the cells to achieve permeabilization of the cell membrane. To avoid any further cells lifting off the glass slides after dehydration with acetone, we added an additional fixation step with 1.6 % PFA to the protocol.

To quench the autofluorescence of neuronal networks, we applied a broad-spectrum LED light on the cells in PBS twice for 45 min. Autofluorescence quenching is another custom adaptation to the standard protocol, loosely based on Du et al., 2019. Before quenching, tissues are usually rehydrated with hydration buffer. However, we noticed a significant amount of cell loss when quenching was conducted after rehydration. Performing quenching prior to incubation with hydration buffer solved this issue.

Incorporating these changes in the protocol enables CODEX multiplex imaging of iPSC-derived neurons without major loss of cells between the staining cycles as shown by brightfield images before and after the procedure (not shown). The stability of our cell cultures during CODEX is further demonstrated in Figure 2. Here we show clusters of nuclei throughout 7 imaging cycles (^~^approximately 21 antibodies). The histogram shows an almost unchanged number of DAPI nuclei throughout the experiment, which confirms that our fixation method was effective. Moreover, and as shown in subsequent sections, we are able to resolve axons and other fine processes throughout our samples, which indicates that even the smallest structures withstood the imaging protocol (Figure 3–5).

### 3.5 Validation of the staining protocol and CODEX panel by imaging iPSC-derived neuronal cultures

To validate, that our protocol allows CODEX multiplex imaging of iPSC-derived neurons, we generated a mixed culture of iPSC-derived cortical, striatal and dopaminergic neurons from the KOLF2.1 control line (Pantazis et al., 2021), which should express all the markers of our human CODEX neuronal phenotyping panel. For the first trial, we tested 32 antibodies already established for CODEX for mouse and human brain tissues, from which 9 (MAP2, NEFL, S100B, SYN, TH, TBR2, CTIP, CAMK2 and CTTN) and the nuclear DAPI staining showed positive signals with expected staining patterns in our iPSC-derived neuronal culture (Supplemental Table 1 and Figure 3A-C). It should be noted that most of the 32 tested antibodies displayed expected staining patterns in other neuronal tissues, including fresh frozen mouse brains and human FFPE brain tissues at Akoya Biosciences (not shown). Functional differences of some antibodies between the different tissue types could be explained by: (a) lack of protein expression depending on the developmental stage of iPSC-derived neurons, (b) higher abundance of the protein in tissues as compared to iPSC-derived neurons, and (c) differential processing of samples (i.e. fixation and quenching) impacts antibody binding. All the listed reasons can result in signals that are below detection level or in unspecific binding of antibodies.

Many of the stained neuronal cells displayed expression of the neuronal markers MAP2, NEFL, and SYN (Figure 3A). As expected, the SYN signal co-localized either with NEFL or MAP2 confirming the specificity of our antibodies. Staining with multiple markers allows for subcategorizing cells. We found expression of TBR2 in a subset of MAP2-positive cells (Figure 3C, cyan arrows) and MAP2-positive cells expressing CTIP2 only (Figure 3C, magenta arrows), or co-expressing both CTIP2 and TBR2 (Figure 3C, white arrows). Cells expressing TBR2 only or TBR2 and CITP2, are most likely deep-layer cortical neurons, while CTIP2 expression only could either indicate cortical or medium spiny neurons. Additional markers (e.g. for striatal GABAergic neurons) would be required to further distinguish between neuronal subtypes. TH-positive cells did not express MAP2 indicating that these were either still immature neurons or neuroendocrine cells. Notably, we further found expected co-localization of proteins known to be involved in synaptogenesis and neurite outgrowth like SYN, CAMK2, and CTTN in neurons (Figure 3B). These patterns were visible in experiments conducted with a 20X lens and were significantly improved during imaging experiments that employed a 40X lens. These results demonstrate that CODEX can be used to study subcellular compartments in iPSC cultures as will be further discussed below.

To further validate, whether the described protocol could be applied to different iPSC lines, we generated induced cortical neurons from an inducible human Ngn2 iPSC line (Wang et al., 2017). Using the CODEX technique, we were able to detect 11 markers (Supplemental Table 2: GFAP, S100B, DCX, SYP, H2A.X, SOX2, MAP2, SYN, NEFL, TH, and EN1) with expected staining patterns (Figure 4A-C). Again, nuclear counterstaining with DAPI as a reference of the total cell number was included. After 45 days of differentiation, we observed many cells positive for the neuronal markers NEFL, MAP2, and SYN (Figure 4A, C). Co-staining for the presynaptic markers SYP and SYN revealed extensive synaptic differentiation and connectivity in our neuronal culture. We also observed puncta without SYN and SYP co-localization (Figure 4, A1, green arrows), potentially indicating immature synapses. Many cells were positive for the neurogenesis marker DCX suggesting that they are either neuronal precursors or young neurons (Karl et al., 2005). Co-localization of DCX and SYP signal in most DCX-positive cells identifies them as young neurons (Figure 4, B1). By staining for the pluripotency marker SOX2 we were able to detect undifferentiated neuronal progenitors (Figure 4, C1, magenta arrows). Importantly, SOX2-positive cells did not co-localize with the mature neuronal marker MAP2 confirming the specificity of our antibodies. We further observed H2A.X-positive cells indicating damaged neurons (Figure 4, C1, red arrows). Even though Ngn2-driven iPSCs are expected to differentiate into cortical neurons, we detected a few TH-positive cells in our culture (Figure 4D). Co-localization with neuronal marker MAP2 further points towards a dopaminergic neuronal identity (Figure 4, D1). This was further confirmed by expression of EN1 in all TH-positive cells (Figure 4, D1, white arrows). Apart from neurons and neuronal precursors, we were able to identify the co-cultured primary astrocytes marked by either S100B, GFAP, or both (Figure 4, B1).

Interestingly, we did not observe positive staining for the cortical marker TBR2 even though the antibody seemed to work in KOLF2.1 iPSC-derived neuronal cultures (Figure 3C). Investigating the gene expression of our human iPSC-derived neuronal cultures by qPCR using SYBR Green array we also did not find upregulated expression of TBR2 in Ngn2-derived cortical neuronal cultures, while we did see upregulation in KOLF2.1-derived neuronal cultures (Supplemental Figure 1). Consequently, the TBR2 protein concentration in the sample was likely below detection level. An explanation for this may be found in the different approaches used for the differentiation process. Forced expression of transcription factors like Ngn2 was found to ‘skip’ steps during neuronal development (Schafer et al., 2019). As TBR2 is a homeobox protein involved in the development of cortical neurons, TBR2 expression might be skipped because of the genetically-induced differentiation process.

Similarly, we did not detect positive SOX2 staining in KOLF2.1-derived neuronal cultures, while we were able to detect SOX2 signal in Ngn2-derived neuronal cultures (Figure 4C). Again, the SYBR Green Array reflects the results seen with antibody staining. At the RNA level, we found upregulated expression of SOX2 in Ngn2-derived neuronal cultures while SOX2 expression was downregulated in KOLF2.1-derived neuronal cultures (Supplemental Figure 1), indicating that the protein SOX2 concentration in the sample was below detection level.

We confirmed the reproducibility of the protocol, by conducting CODEX multiplex-imaging of iPSC-derived neurons both at laboratories at Stanford Medicine and at Akoya Biosciences as shown in Figures 3, 4, and Supplemental Figure 2, respectively. Each experiment employed different instrumentation and operators. Additional antibodies were included in separate experiments to validate their functionality and to expand our CODEX panel for iPSC-derived neurons (Supplemental Figure 2 and Figure 5).

### 3.6 Improving the readout of CODEX imaging by increasing the magnification of the microscope

One potential disadvantage of the CODEX technique is that imaging approaches use only low magnification lenses. While this may be optimal to study large tissue samples, deep phenotyping of iPSC-derived neurons, especially for analyzing neuronal network connectivity or synaptic phenotypes, will require higher magnifications. Consequently, we tested whether we could conduct CODEX multiplex-imaging of iPSC-derived neurons using a 40x oil immersion lens. We were able to re-image five different markers including MAP2, the voltage-gated R-type calcium channel alpha subunit (CAV2.3), the dopamine- and cAMP-regulated protein phosphatase 1 regulatory inhibitor subunit 1B (DARPP32), the neuronal protein alpha-synuclein (aSyn), and the presynaptic active zone protein piccolo (PCLO) in a sample that was already imaged with a 20x objective showing the robustness of the adapted fixation protocol (Figure 5: image with 40x objective, Supplemental Figure 3: image with 20x objective). Both CAV2.3 (Dietrich et al., 2003; Heinrich and Ryglewski, 2020), DARPP32 (Svenningsson et al., 2004), aSyn (Burré, 2015), and PCLO (Gundelfinger et al., 2016) are known modulators of synaptic neurotransmission. Therefore, it is not surprising that we found co-localization of these proteins in dendritic spines (dendrites are marked by MAP2 staining) and synaptic terminals (Figure 5A, B; white arrows). Our data are unique in showing successful CODEX imaging with subcellular resolution, using a 40x oil-immersion lens. This approach opens up the possibility of using even higher magnification lenses (e.g. 60x) for future applications. This will enable better analysis of biomarkers throughout the substructure of individual cells.

## 4. Discussion

Generation of iPSC-derived neurons is labor-intensive, costly, lengthy, and variable depending on the experiment and clonal lines. Hence, maximizing the readout of iPSC-derived neural networks through multiplex imaging will be critical to improving the input-output ratio. The biggest limitation in staining and imaging neuronal cultures is the risk of losing cells even during standard staining procedures. Therefore, multiplex imaging applications that utilize multiple washing and imaging cycles prove to be especially challenging and demand a specialized preparation protocol for iPSC-derived neurons. Here we report multiplex imaging with CODEX for *in vitro* differentiated neurons from human iPSCs while overcoming that challenge. We developed an optimized CODEX panel for (1) neuronal phenotyping (Figure 6), (2) histological preparation to processing fragile neuronal networks, (3) reduction of autofluorescence of human neuronal tissues, and (4) CODEX imaging with improved resolution (higher magnification objectives).

**Figure 6:**
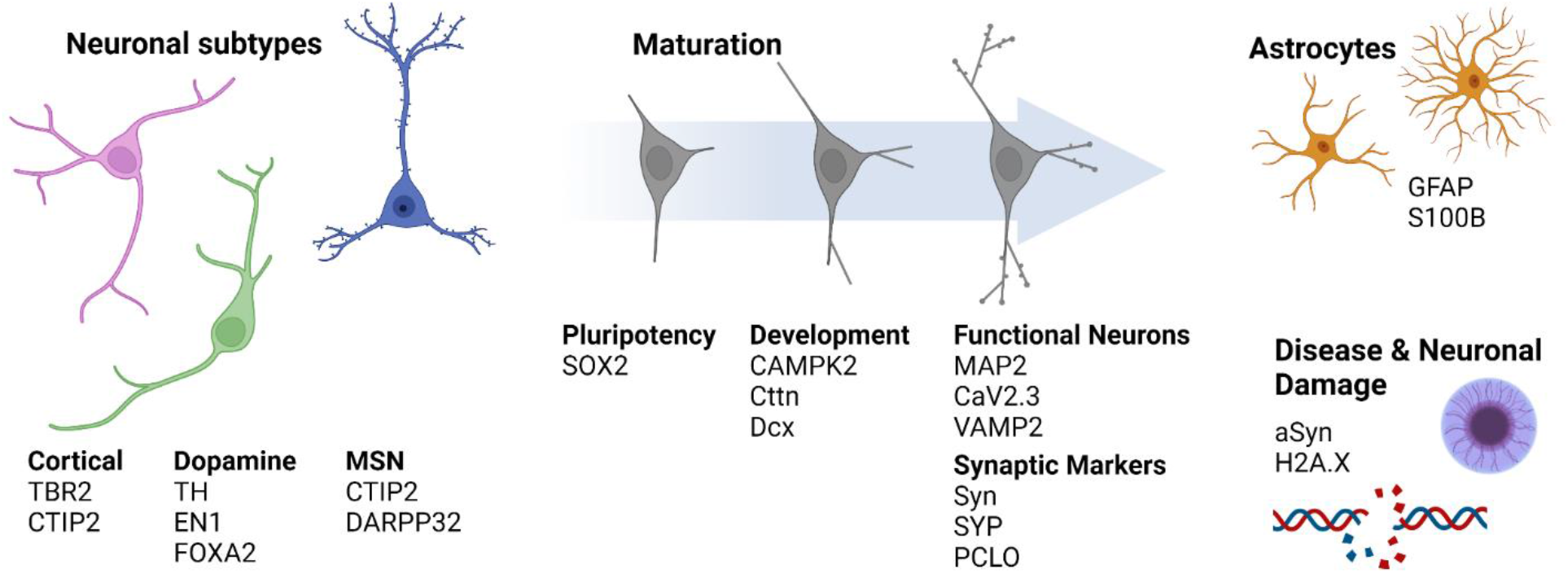
CODEX phenotyping panel for iPSC-derived neuronal cultures. Antibody panel for characterizing iPSC-derived neurons using CODEX multiplex imaging. Markers are aimed at neuronal subtypes like cortical T-box brain protein 2 (TBR2), chicken ovalbumin upstream promoter-transcription factor-interacting protein 2 (CTIP2), dopaminergic neurons (TH, EN1, FOXA2), and medium spiny neurons (CTIP2, DARPP32); markers for pluripotency (SOX2), neurodevelopment (CAMPK2, CTTN, DCX) and mature neurons (MAP2, CAV2.3, VAMP2) displaying synaptic markers (SYN, SYP, PCLO); markers for astrocytes (GFAP, (S100B); as well as disease markers (alpha-synuclein (aSyn)) and markers for neuronal damage (H2A.X).

The major advantage of CODEX fluorophores is that they circumvent the primary antibody host interaction with oligonucleotide barcodes that allow for more flexible, yet specific primary antibody binding. However, this also means that the primary antibodies need to be conjugated to the complementary DNA oligonucleotides. The conjugation procedure requires carrier-free antibodies, which currently limits the choice of many commercial antibodies. Staining samples with multiple antibodies further increases the risk of cross-reactivity between antigens and/or oligonucleotides, thus antibodies that work well under standard immunocytochemistry conditions might not work for CODEX or result in unexpected staining patterns. Consequently, building a CODEX panel can be expensive and time-consuming, especially when starting with new samples such as iPSC-derived neurons that involve a prolonged testing and optimization process. However, once optimized for cell types of interest, the potential for imaging multiple markers in one sample greatly exceeds that of traditional immunofluorescence. To verify that the antibodies are truly binding to proteins being expressed in the differentiated neurons, it is necessary to incorporate a complementary technique in this optimization process such as SYBR Green Q-PCR expression analysis which we successfully used for our differentiated neurons to verify expression at mRNA level. Additionally, optimization of antibodies is crucial to make sure there is no cross-talk between different barcode-conjugated antibodies and to achieve the best signal to noise during the imaging process. During this study, we tested 38 antibodies in three different samples and were able to optimize 21 in our CODEX panel for iPSC-derived neurons (Supplementary Tables 1-4). The other 19 antibodies that were used in the CODEX panel either had unspecific binding or the protein was not yet expressed in the iPSC-derived neuronal population that was tested here (Supplementary Table 5).

CODEX imaging of iPSC-derived neurons will need additional optimization and refinement to improve image quality. We identified three areas that need further improvement: (1) sample thickness, (2) raw data processing, and (3) resolution. First, for optimal imaging conditions with CODEX, tissues sections should be 5-8μm thick. However, during the differentiation process, the cell bodies of iPSC-derived neuronal networks tend to grow in multiple layers that exceed the maximum imaging thickness. Optimizing the culturing conditions or clearing iPSC-derived neuronal cultures to achieve monolayer cultures would improve the imaging quality with CODEX. Second, the processing and segmenting of raw images is a challenge. Available segmentation algorithms operate on the assumption of circular cells or nuclei, hence the development of new software or optimization of segmentation algorithms will be critical for processing images of irregularly shaped cells, such as neuronal cell cultures. Finally, current hardware limitations (microscope and lenses) only enable imaging of samples at low magnification. To use the CODEX panel for deep phenotyping of iPSC-derived patient neurons compared to controls in future applications, higher resolutions will be needed, especially for investigating the network connectivity or synaptic phenotypes.

In this study, we report for the first time CODEX multiplex imaging with a 40x oil-immersion objective, which currently requires manual application of oil during the imaging procedure, limiting the imaging time, number of multiplexed markers, and size of the imaging area. Although these disadvantages will become more pronounced with higher magnification lenses, we expect this to improve with future hardware and software advancements in the field.

The CODEX technology in human iPSC neurons permits a high level of multiplexing which has not been possible with current immunofluorescent imaging techniques. Future versions of this human CODEX panel will expand on markers for cell types including microglia (e.g. CD11b, CD45), additional markers for neuronal subtypes including markers for cortical neurons (e.g. POU domain, class 3, transcription factor 2 (BRN2), vesicular glutamate transporter 1 (VGLUT1), or DNA-binding protein SATB2), striatal neurons (e.g. Forkhead box protein P1 (FOXP1), dopamine receptor DRD1/2, or vesicular GABA transporter (VGAT)), and dopaminergic neurons (e.g. transcription factors NURR1 and LMX1A), as well as further markers for connectivity including post-synaptic markers like postsynaptic density protein PSD95, and disease-specific markers (e.g. TAU, PARKIN, P62). This panel will be invaluable for increasing the input-output ratio of iPSC-derived neuronal cultures and deep phenotyping of iPSC-derived neurons leading to a better understanding of iPSC-based model systems.

## 5. Conclusion

We applied and adapted CODEX multiplex-imaging to analyze human iPSC-derived neuronal cultures of cortical, striatal, and dopaminergic neurons. We developed and optimized multiple CODEX antibody and reporter panels, and validated these CODEX panels against conventional immunofluorescence or mRNA expression of tested proteins. Our approach presents an innovative new means for the characterization of iPSC-derived neuronal cultures, discovery of cell-type composition, and developmental differences in human iPSC neuronal cultures. As such, this technique could become an important vehicle for protocol development and quality control, molecular characterization, and characterization of disease-specific differences in *in vitro* cultures that were previously unrecognized due to the lack of a highly multiplexed imaging methodologies.

## Abbreviations

iPSC: induced pluripotent stem cells
CODEX: CO-Detection by indexing
DIV: days in vitro
ICC: immunocytochemistry
NPCs: neural progenitor cells
KO-DMEM: knockout Dulbecco’s modified Eagle’s medium
PBS: phosphate-buffered saline
THZ: Thiazovivin
RT: room temperature
LGE: lateral ganglionic eminence
MSN: medium-sized spiny neurons
KSR: Knockout serum replacement
PenStrep: penicillin, streptomycin
NEAA: non-essential amino acids
Vit. A: vitamin A
FBS: fetal bovine serum
Ngn2: neurogenin 2
NT3: neurotrophin 3
cDNA: complementary DNA
NC: negative control
ddH2O: double-distilled water
PFA: paraformaldehyde
MAV: Multiplex Analysis Viewer
BSA: bovine serum albumin
PLL: poly-L-lysine
BS3: bis(sulfosuccinimidyl)suberate

## 8. Consent for publication

Not applicable.

## 9. Ethical approval and consent to participate

Not applicable. Human cell line KOLF2.1 was obtained from Jackson Laboratories. Human cell line WTC11-G3 was obtained from Gladstone Institutes.

## 12. Authors contributions

BS, OB designed the study and oversaw all aspects of the study. LH, FZ, AT prepared the human iPSC neurons. JS, CH, NN, AK, JM performed CODEX experiments. AT, JS validated antibodies for CODEX. MYC performed SYBR Green experiments. LH and FZ wrote the first draft of the manuscript. All co-authors interpreted and analyzed the data and critically revised the manuscript.

## Acknowledgments

We thank Abcam, Cell Signaling Technologies (CST), and Synaptic Systems for selecting and providing suitable antibody candidates for this study.

