## Supplementary Materials for "Multiplex imaging of human induced pluripotent stem cell-derived neurons with CO-Detection by indEXing (CODEX) technology"

Department Pathology

Stanford University School of Medicine

300 Pasteur Dr, R271/217

Stanford, CA 94305

### Supplemental Table 1: Antibody Panel for iPSC-derived Neurons for Figure 2

| Antibody | Cell type-specific | AB clone | Vendor | Cat. No. | Barcode | Dilution | Exposure (ms) | Fluoro-phore |
|---|---|---|---|---|---|---|---|---|
| S100B | Astrocyte | RM304 | RevMab | | BX032 | 1:50 | 350ms | 550 |
| GFAP | Astrocyte | 134B1 | Synaptic Systems | 173011 | BX007 | 1:50 | 350ms | Cy5 |
| TBR2 | Cortical layer transcriptional activator | EPR21950-241 | Abcam | ab261913 | BX002 | 1:50 | 350ms | 550 |
| CTIP2 | Cortical Layer V, Medium Spiny Neurons | EPR23120-25 | Abcam | ab269367 | BX043 | 1:50 | 250ms | 488 |
| Tyrosine Hydroxylase | Dopaminergic neuron | E2L6M | Biolegend | 818001 | BX049 | 1:50 | 350ms | Cy5 |
| Neurofilament-L (NEFL) | Mature neuronal cells | C28E10 | Cell Signaling Tech | 2837BF | BX022 | 1:50 | 250ms | 488 |
| MAP2ab | Mature neuronal cells | MT-08 | Antibodies Online.com | ABIN125739 | BX023 | 1:50 | 350ms | 550 |
| Synapsin | Synaptic protein | 7H10G6 | Antibodies Online.com | ABIN5542390 | BX001 | 1:50 | 250ms | 488 |
| CAMK2 | 100% CaMK2 beta, 93% CaMK2 gamma and 73% CaMK2 alpha and delta, cytosol, synapse and dendrites | EPR6686(2) | Abcam | ab227108 | BX004 | 1:50 | 350ms | Cy5 |

### Supplemental Table 2: Antibody Panel for iPSC-derived Neurons for Figure 3

| Antibody | Cell type-specific | AB clone | Vendor | Cat. No. | Barcode | Dilution | Exposure (ms) | Fluoro-phore |
|---|---|---|---|---|---|---|---|---|
| S100B | Astrocyte | RM304 | RevMab | 31-1189-00 | BX036 | 1:50 | 350ms | 550 |
| GFAP | Astrocyte | 134B1 | Synaptic Systems | 173011 | BX007 | 1:50 | 350ms | Cy5 |
| Tyrosine Hydroxylase | Dopaminergic neurons | E2L6M | Biolegend | 818001 | BX049 | 1:50 | 250ms | 488 |
| Engrail1 (EN1) | Ectodermal Transcription factor | 1F5 | Thermo Fisher | H00002019-M06 | BX041 | 1:50 | 350ms | 550 |
| H2A.x | Histone H2A, DNA repair | 2F3 | Biolegend | 613402 | BX042 | 1:50 | 350ms | Cy5 |
| Neurofilament-L (NEL) | Mature neuronal cells | C28E10 | Cell Signaling Tech | 2837BF | BX022 | 1:50 | 250ms | 488 |
| MAP2ab | Mature neuronal cells | MT-08 | Antibodies Online.com | ABIN125739 | BX023 | 1:50 | 350ms | 550 |
| SOX2 | Neuroprogenitor | EPR3131 | Abcam | ab215970 | BX045 | 1:50 | 350ms | Cy5 |
| Synaptophysin (SYP) | Presynaptic vesicles of neurons | Sy38 | Invitrogen/ Thermo | MA1-213 | BX028 | 1:50 | 250ms | 488 |
| Synapsin 1 (SYN) | Synaptic protein | 7H10G6 | Antibodies Online.com | ABIN5542390 | BX001 | 1:50 | 250ms | 488 |
| Doublecortin (DCX) | Mature neuronal cells | EPR19997 | Abcam | ab222921 | BX034 | 1:50 | 250ms | 488 |

**Supplemental Table 3: Antibody Panel for iPSC-derived Neurons for Figure 4**

| Antibody | Cell type-specific | AB clone | Vendor | Cat. No. | Barcode | Dilution | Exposure (ms) | Fluoro-phore |
|---|---|---|---|---|---|---|---|---|
| MAP2ab | Mature neuronal cells | MT-08 | Antibodies Online.com | ABIN125739 | BX023 | 1:50 | 350ms | 550 |
| α Synuclein | Synaptic protein | 354A10 | Synaptic Systems | 128211 | BX004 | 1:50 | 250ms | 488 |
| CAV2.3 | Voltage gated calcium channel | 62C10 | Synaptic Systems | 152403 | BX042 | 1:50 | 350ms | Cy5 |
| Piccolo | Synaptic active zone marker | Neu 287 | Synaptic Systems | 142003 | BX003 | 1:50 | 350ms | Cy5 |
| DARPP32 | Marker for striatal medium spiny neurons | polyclonal | Abcam | 382003 | BX024 | 1:50 | 350ms | Cy5 |

**Supplemental Table 4: Antibody Panel for iPSC-derived Neurons for Supplemental Figure 2**

| Antibody | Cell type-specific | AB clone | Vendor | Cat. No. | Barcode | Dilution | Exposure (ms) | Fluoro-phore |
|---|---|---|---|---|---|---|---|---|
| Neurofilament-L (NEFL) | Mature neuronal cells | C28E10 | Cell Signaling Tech | 2837BF | BX022 | 1:50 | 250ms | 488 |
| Synaptophysin (SYP) | Presynaptic vesicles of neuron | Sy38 | Invitrogen / Thermo | MA1-213 | BX028 | 1:50 | 250ms | 488 |
| Tyrosine Hydroxylase (TH) | Dopaminergic neuron | polyclonal | Synaptic Systems | 213104 | BX049 | 1:50 | 250ms | 488 |
| α Synuclein | Neuronal protein | 354A10 | Synaptic Systems | 128211 | BX004 | 1:50 | 250ms | 488 |
| FOXA2 | Floorplate marker expressed in Dopaminergic neuron | 7E6 | Abcam | ab60721 | BX036 | 1:50 | 350ms | Cy5 |
| VAMP2 | Vesicle protein, Synaptic marker | 69.1 | Synaptic Systems | 104008 | BX010 | 1:50 | 250ms | 488 |

**Supplemental Table 5: Other antibodies tested against iPSCs (failed or no staining observed in iPSC culture samples)**

| Antibody | Cell type-specific | AB clone | Vendor | Cat. No. | Barcode | Dilution | Exposure | Fluorophore |
|---|---|---|---|---|---|---|---|---|
| VGlut1 | Glutamate transporter of synaptic vesicles | EPR22269 | Abcam | ab242017 | BX050 | 1:50 | 250ms | 488 |
| GAD67 | Catalyzes the production of GABA | EPR20578 | Abcam | ab246335 | BX029 | 1:50 | 350ms | 550 |
| DAT | Transmembrane protein to Reuptake of dopamine | EP19695 | Abcam | ab221845 | BX025 | 1:50 | 350ms | Cy5 |
| MAG | Axon-myelin stabilization | EPR24276-125 | Abcam | ab277535 | BX024 | 1:50 | 350ms | Cy5 |
| Serotonin transporter | Regulates serotonergic neurotransmission | EPR23530-3 | Abcam | ab275094 | BX013 | 1:50 | 250ms | 488 |
| TUBB3 | Mature neuronal cells | Tu-20 | Antibodies Online.com | ABIN93911 | BX010 | 1:50 | 350ms | 550 |
| ACTIN | Cytoskeletal protein | S12-I | Antibodies Online.com | ABIN870295 | BX006 | 1:50 | 350ms | Cy5 |
| SLC32 | Uptake of GABA into Synaptic vesicles | SLC32A1 | Antibodies Online.com | ABIN2855225 | BX031 | 1:50 | 250ms | 488 |
| Cortactin | Regulate Ca leveling in neurons | Polyclonal | Antibodies Online.com | ABIN2854674 | BX014 | 1:50 | 350ms | 550 |
| TMEM119 | Microglia | E3E4T | Cell Signalling Tech | 1134BF | BX027 | 1:50 | 350ms | Cy5 |
| Basson | Synaptic protein | Polyclonal | Antibodies Online.com | ABIN863198 | BX019 | 1:50 | 250ms | 488 |
| Shank3 | Synaptic protein | S367-51 | Antibodies Online.com | ABIN1741251 | BX020 | 1:50 | 350ms | 550 |
| SaTB | Cortical Layer IV | EPNCIR130A | Abcam | ab212177 | BX047 | 1:50 | 350ms | 550 |
| Calbindin | Calcium-binding protein | EP3478 | Abcam | ab233018 | BX046 | 1:50 | 350ms | Cy5 |
| PSD-95 | Synaptic protein | 6G6 | Antibodies Online.com | ABIN361694 | BX026 | 1:50 | 350ms | 550 |
| TAU | Mature neuronal cells | EPR22524-94 | Abcam | ab255271 | BX003 | 1:50 | 350ms | Cy5 |
| GRIN2B | Myelin sheath of oligodendrocytes and Schwann cells | 6E9A8 | Antibodies Online.com | ABIN5611338 | BX016 | 1:50 | 250ms | 488 |
| SLC17 | Glutamate transporter at SLC family | S28-9 | Antibodies Online.com | ABIN1027710 | BX017 | 1:50 | 350ms | 550 |
| MBP | Myelin sheath of oligodendrocytes and Schwann cells | EPR21188 | Abcam | ab218011 | BX037 | 1:50 | 250ms | 488 |
| CD34 | Endothelium | Qbend10 | Invitrogen/ Thermo | MA1-10202 | BX005 | 1:50 | 350ms | 550 |
| Nestin | Pre-neuronal | 10C2 | Biolegend | 656802 | BX035 | 1:50 | 350ms | 550 |

**Supplemental Table 6: Materials and Reagents**

| Item name | Vendor | Catalog # |
|---|---|---|
| **iPSC maintenance:** | | |
| Matrigel | Thermo Fisher Scientific | CB-40230 |
| KnockOut DMEM | Thermo Fisher Scientific | 10829018 |
| Bambanker | Thermo Fisher Scientific | NC9582225 |
| Stemflex media kit | Thermo Fisher Scientific | A3349401 |
| Thiazovivin | ReproCell | 04-0017 |
| ReLeSR | StemCell Technologies | 05872 |
| **Neuronal differentiation of iPSC** | | |
| KnockOut™ Serum Replacement | Thermo Fisher Scientific | 10828028 |
| Penicillin Streptomycin | Thermo Fisher Scientific | 15140-122 |
| GlutaMAX | Thermo Fisher Scientific | 35050-061 |
| NEAA | Thermo Fisher Scientific | 11140050 |
| b-Mercaptoethanol | Thermo Fisher Scientific | 21985023 |
| Knockout DMEM/F12 | Thermo Fisher Scientific | 12660-012 |
| B27 Supplement 50X without Vit.A | Thermo Fisher Scientific | 12587010 |
| Bd-cAMP | Peprotech | 6099240 |
| L-Ascorbic Acid | Peprotech | 5088177 |
| DAPT | Peprotech | 2634 |
| Neurobasal A Medium | Thermo Fisher Scientific | 12349015 |
| BDNF | Peprotech | 450-02 |
| GDNF | Peprotech | 450-10 |
| FBS | Thermo Fisher Scientific | SH3091003 |
| N2 Supplement 100X | Thermo Fisher Scientific | 17502-048 |
| Neurobasal™ Medium | Thermo Fisher Scientific | 21103049 |
| DMEM/F-12, no glutamine | Thermo Fisher Scientific | 21331020 |
| neurotrophin-3 | Peprotech | 450-03 |
| Doxycycline hyclate | Sigma-Aldrich | D9891-5G |
| Cultureone™ Supplement (100X) | Thermo Fisher Scientific | A3320201 |
| **Gene expression using SYBR green** | | |
| 2-mercaptoethanol | Sigma Aldrich | M2650 |
| DNase I, Amplification Grade | Thermo Fisher Scientific | 18068015 |
| Ethanol, Molecular grade | Thermo Fisher Scientific | BP2818500 |
| Homogenizer spin column | Thermo Fisher Scientific | 12183-026 |
| PureLink RNA Mini Kit | Thermo Fisher Scientific | 12183025 |
| RNase Away | Thermo Fisher Scientific | 10328011 |
| High-Capacity cDNA Reverse Transcription Kit | Thermo Fisher Scientific | 4368814 |
| MicroAmp 8 tube strip | Thermo Fisher Scientific | A30589 |
| 384-well PCR plate | Thermo Fisher Scientific | AB1384W |
| Nuclease-free water | US Biological LifeSciences | W0900 |
| PowerUp™ SYBR™ Green master mix | Thermo Fisher Scientific | A25780 |
| **CODEX coverslip preparation and NPCs seeding** | | |
| Coverslips | Electron Microscopy Science | 72204-10 |
| Poly-L-Lysine | Sigma-Aldrich | P8920 |
| Laminin (mouse) | Thermo Fisher Scientific | 23017-015 |
| **Fixation, hydration, and Pre-staining fixation of neurons of coverslip** | | |
| PFA | Thermo Fisher Scientific | 50-980-495 |
| Acetone 100% | Millipore-Sigma | AX0120-6 |

| | | |
|---|---|---|
| Drierite absorbent beads | W.A. Hammond Drierite Co | 21001 |
| Hydration Buffer | Akoya Biosciences | 7000008 |
| Staining buffer | Akoya Biosciences | 7000008 |
| Storage buffer | Akoya Biosciences | 7000008 |
| Blocking Buffer | Akoya Biosciences | 7000008 |
| PBS | MP Biomedicals | 091860454 |
| Methanol | Sigma-Aldrich | 646377 |
| BS3 substrate | Thermo Fisher Scientific | 21580 |
| **Primary Antibody processing** | | |
| CODEX conjugation kit | Akoya Biosciences | 7000009 |
| BSA removal kit | Abcam | Ab173231 |
| **Primary Astrocyte culture** | | |
| Human Astrocytes | ScienceCell | 1800 |
| Ara-C | Sigma-Aldrich | C6645 |
| Astrocyte Media | ScienceCell | 1801 |
| **Barcodes** | | |
| BX001 | Akoya Biosciences | 5450013 |
| BX002 | Akoya Biosciences | 5450023 |
| BX003 | Akoya Biosciences | 5450026 |
| BX004 | Akoya Biosciences | 5450014 |
| BX005 | Akoya Biosciences | 5450024 |
| BX006 | Akoya Biosciences | 5450027 |
| BX007 | Akoya Biosciences | 5450015 |
| BX010 | Akoya Biosciences | 5450016 |
| BX013 | Akoya Biosciences | 5450017 |
| BX014 | Akoya Biosciences | 5450025 |
| BX016 | Akoya Biosciences | 5150001 |
| BX017 | Akoya Biosciences | 5250001 |
| BX019 | Akoya Biosciences | 5150002 |
| BX020 | Akoya Biosciences | 5250002 |
| BX022 | Akoya Biosciences | 5150003 |
| BX023 | Akoya Biosciences | 5250003 |
| BX024 | Akoya Biosciences | 5350003 |
| BX025 | Akoya Biosciences | 5150004 |
| BX026 | Akoya Biosciences | 5250004 |
| BX027 | Akoya Biosciences | 5350004 |
| BX028 | Akoya Biosciences | 5150005 |
| BX029 | Akoya Biosciences | 5250005 |
| BX031 | Akoya Biosciences | 5150006 |
| BX032 | Akoya Biosciences | 5250006 |
| BX035 | Akoya Biosciences | 5250007 |
| BX036 | Akoya Biosciences | 5350007 |
| BX037 | Akoya Biosciences | 5150008 |
| BX041 | Akoya Biosciences | 5250008 |
| BX042 | Akoya Biosciences | 5350008 |
| BX043 | Akoya Biosciences | 5150010 |
| BX045 | Akoya Biosciences | 5350009 |
| BX046 | Akoya Biosciences | 5150011 |
| BX047 | Akoya Biosciences | 5250009 |
| BX049 | Akoya Biosciences | 5150012 |

| | | |
|---|---|---|
| BX050 | Akoya Biosciences | provided by Akoya |

**Supplemental Table 7: Primers used for SYBR Green qPCR assay**

| Gene | Function | Primer direction (5' to 3') | Sequence |
|---|---|---|---|
| FOXA2 | Dopaminergic neuron marker | Forward | CTGGGAGCGGTGAAGATGGAA |
| | | Reverse | TTCATGTTGCTCACGGAGGAGTAG |
| EN1 | Dopaminergic neuron marker | Forward | TGCTAGATAAGAACGAGCGATCCA |
| | | Reverse | GAGGAAGGAGGCAGGCGAAG |
| Doublecortin | Neurodevelopmental marker | Forward | TCTGACAACATCAACCTGCCTCA |
| | | Reverse | TTCCTCCAGTTCATCCATGCTTCC |
| LMX1A | Dopaminergic neuron marker | Forward | CTTCTGCTGCTGTGTCTGCG |
| | | Reverse | CTCCCGCTCCTTCTCATAGTCC |
| SOX2 | Pluripotency marker | Forward | TACATGAACGGCTCGCCCAC |
| | | Reverse | GGACTTGACCACCGAACCCA |
| MAP2 | Neuronal maturation marker | Forward | GGACATGATCTTTCTCCTCTGGCTT |
| | | Reverse | AGGTGTGGTGGCTGGAAGGT |
| TH | Dopaminergic neuron marker | Forward | ATTGCTGAGATCGCCTTCCAGTA |
| | | Reverse | GTGGTGTAGACCTCCTTCCAGG |
| GFAP | Astrocyte marker | Forward | GGGACAATCTGGCACAGGAC |
| | | Reverse | GGGTGGCTTCATCTGCTTCC |
| S100B | Astrocyte marker | Forward | ATATTCTGGAAGGGAGGGAGACAA |
| | | Reverse | AGAAATGGGAAAGCTCATTGTTGATG |
| H2A.X | Cell stress and damage | Forward | GCAGTGCTGGAGTACCTCACC |
| | | Reverse | CTCCTCGTCGTTGCGGATGG |
| CAMKII | Neurodevelopmental marker | Forward | CAGACTTCGGCCTAGCTATCGAG |
| | | Reverse | GGCTTGCCATACGCCTCTTTG |
| CTIP2 | Striatal & Cortical neuron marker | Forward | ATTTCCTGGGCGACAGCAACC |
| | | Reverse | GACTGAAGAGAGGCGGCGTG |
| Synapsin | Synaptic marker | Forward | GGTGAAGGTCGTGCGGTCTC |
| | | Reverse | GTAGTCTCCGTTGCGTGCCAT |
| Synaptophysin | Synaptic marker | Forward | TCCCTTTCCCTGCATCCCTTG |
| | | Reverse | ACACAGCCGAGGTCTGTTCC |
| TBR2 | Cortical Development | Forward | AATGTGTTCGTAGAGGTGGTGCT |
| | | Reverse | CCCTGCATGTTATTGTCGGCTTT |
| SNCA | Presynaptic protein, Disease marker for Parkinson's | Forward | GCAGAAGCAGCAGGAAAGACAAA |
| | | Reverse | CACTTGCTCTTTGGTCTTCTCAGC |
| Cortactin | Neurodevelopmental marker | Forward | TGTCTTTCAAGAGCATCAGACCCT |
| | | Reverse | CCACACCAAATTTCCCTCCATAGC |

| | | | |
|---|---|---|---|
| MAPT | Neuronal maturation marker | Forward | AGTCGAAGATTGGGTCCCTGG |
| | | Reverse | GGAAGGTCAGCTTGTGGGTTTC |
| GAD65 | Striatal/GABA-ergic marker | Forward | TTGGATATGGTTGGATTAGCAGCAG |
| | | Reverse | GCACAAATACTGGAGCAATTTCATAGG |
| VGAT | Striatal/GABA-ergic marker | Forward | GAGCAAGCGGAGATAGCGACTTT |
| | | Reverse | GGACAGCGGAAAGGACAGAAGG |
| Neurofilament | Neuronal marker | Forward | CAAGACCTCCTCAACGTGAAGATG |
| | | Reverse | TGAAACTGAGTCGGGTCTCCTC |
| GAPDH | Housekeeping Gene | Forward | AAGGCTGTGGGCAAGGTCATC |
| | | Reverse | GGCAGGTCAGGTCCACCACT |

**Supplemental Table 8: Thermal cycling settings for qPCR reaction**

| Step | Temperature | Duration | Cycles |
|---|---|---|---|
| UDG activation | 50 °C | 2 min | Hold |
| Dual-Lock™ Taq DNA polymerase | 95 °C | 2 min | Hold |
| Denature | 95 °C | 15 s | 40 |
| Anneal/ extend | 60 °C | 1 min | |

**Supplemental Table 9: Settings for qPCR dissociation step**

| Step | Ramp rate | Temperature | Time |
|---|---|---|---|
| Denature | 1.6 °C/s | 95 °C | 15 s |
| Anneal | 1.6 °C/ s | 60 °C | 1 min |
| Dissociation | 0.15 °C/ s | 95 °C | 15 s |

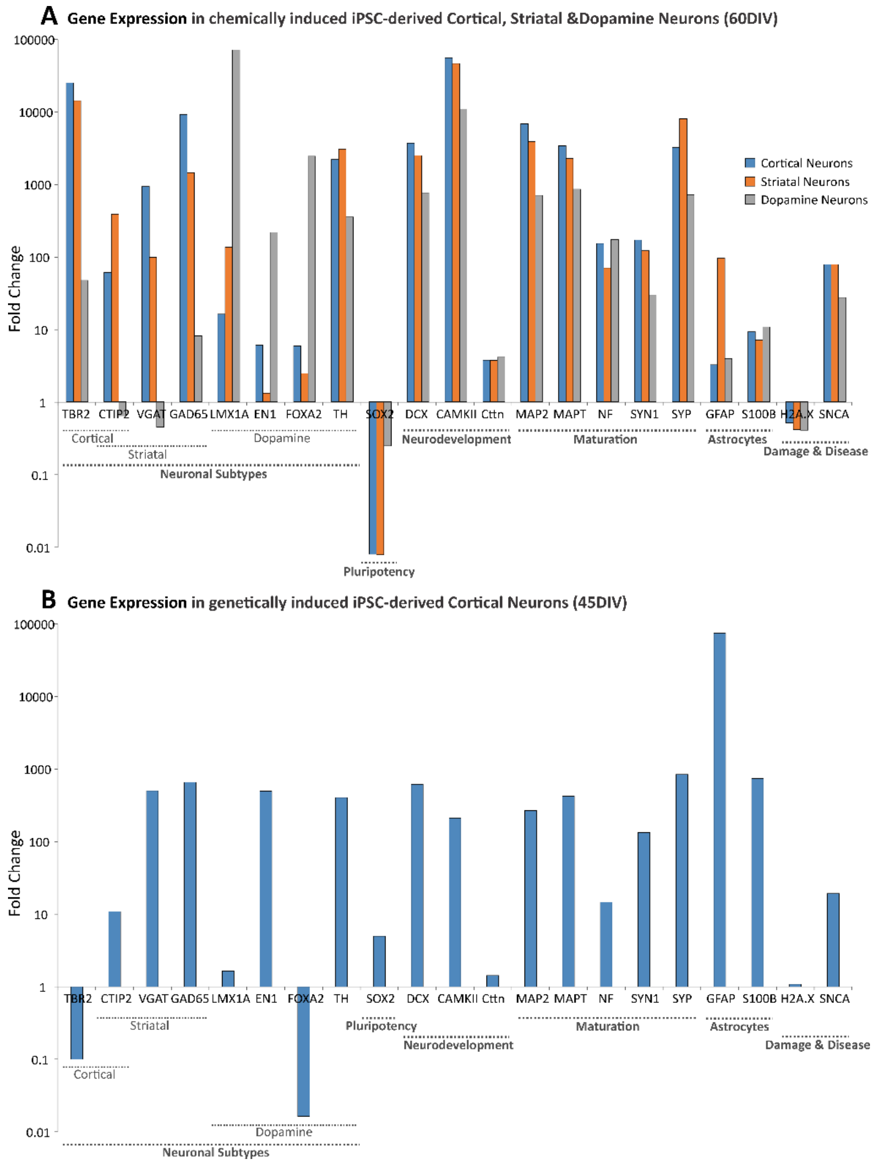


***Supplemental Figure 1: Analyzing the gene expression of human iPSC-derived neurons using the SybrGreen assay.*** *Gene expression was analyzed in human induced pluripotent stem cells (iPSCs)-derived neurons and the undifferentiated iPSC line (reference sample; (**A**) KOLF2.1 or (**B**) Neurogenin 2 (WTC11_G3) iPSC line) by qPCR using the SYBR Green assay for the 21 genes of the CODEX panel.Expression of glyceraldehyde-3-phosphate dehydrogenase (GADPH) was measured as a loading control. Per gene, the cycle threshold (Ct) value was measured and ΔCt was calculated by normalizing the Ct value of the target gene to the Ct value of the loading control (ΔCt = Ct of the target – Ct of the reference). Then ΔΔCT was calculated by normalizing ΔCt of the target gene in our neuronal samples to ΔCt of the same gene in the undifferentiated iPSC line (ΔΔCT = ΔCT(a target sample)−ΔCT(a reference sample)). Finally, the fold change was calculated (fold change =* $2^{-\Delta\Delta CT}$*). [**A**] Human KOLF2.1 iPSCs were differentiated into cortical (blue), GABAergic striatal (orange), or dopaminergic (grey) neurons using chemical induction. After 60 days in vitro (DIV), qPCR by SYBR Green assay was conducted to analyze gene expression. Gene expression was also measured in the undifferentiated KOLF2.1 iPSC line to calculate the fold change. [**B**] Inducible human Ngn2 iPSCs were differentiated into induced cortical neurons. After 45 DIV qPCR by SYBR Green assay was conducted to analyze gene expression. Gene expression was also measured in the undifferentiated Ngn2 iPSC line to calculate the fold change.*

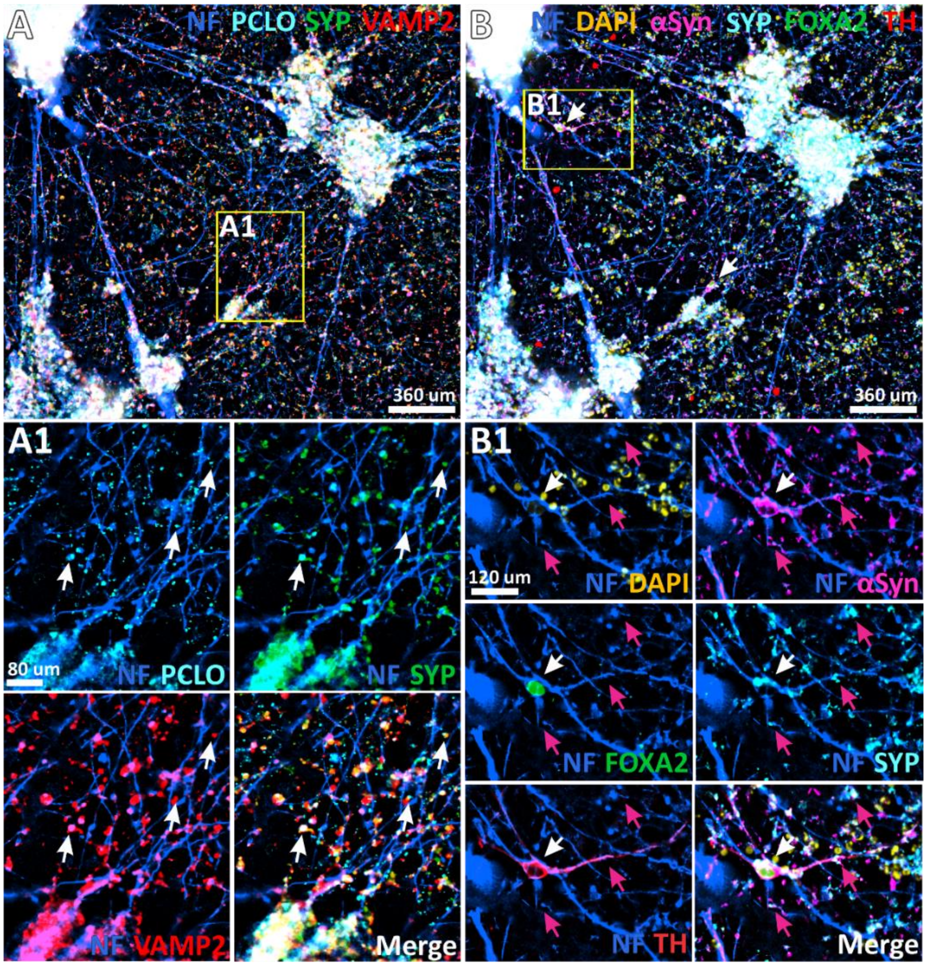


***Supplemental Figure 2: CODEX imaging of iPSC-derived neurons (70 DIV).*** *Human KOLF2.1 iPSCs were differentiated into a mixed culture of cortical, striatal, and dopaminergic neurons. Codex imaging is shown for 7 different markers. Scale bars represent either 360µm, 120µm, or 80µm. [**A**] Representative image of Neurofilament (NEFL, blue), piccolo (PCLO, cyan), Synaptophysin (SYP, green), and the vesicle-associated membrane protein 2 (VAMP2, red) staining. [**A1**] Magnification of the selected area (yellow square) highlighting PCLO (cyan), SYP (green), and VAMP2 (red) colocalization (white arrows) in NEFL-positive cells. [**B**] Representative image of Neurofilament (NF, blue), DAPI (yellow), alpha-synuclein (aSyn, magenta), forkhead-box-protein A2 (FOXA2, green), SYP (cyan), and tyrosine hydroxylase (TH, red) staining. [**B1**] Magnification of selected area (yellow square) highlighting FOXA2 (green) and TH (red) colocalization (white arrows) in NEFL labeled neurons. DAPI staining was higher in potentially damaged cells (bright yellow dots) and was therefore detectable but difficult to see in nuclei of healthy neurons (e.g. FOXA2-positive nuclei).*
